# Displacement and dissociation of oligonucleotides during DNA hairpin closure under strain

**DOI:** 10.1101/2022.04.30.490171

**Authors:** Fangyuan Ding, Simona Cocco, Saurabh Raj, Maria Manosas, Michelle M. Spiering, David Bensimon, Jean-François Allemand, Vincent Croquette

## Abstract

The simple hybridization kinetic of an oligonucleotide to its template is a fundamental step in many biological processes such as replication arrest, CRISPR recognition, DNA sequencing, DNA origami, etc. Although approaches exist that address special cases of this problem, there are no simple general prediction schemes. In this work, we have measured experimentally, with no fluorescent labelling, the displacement of an oligonucleotide from its substrate in two situations: one corresponding to oligonucleotide binding/unbinding on ssDNA and one where the oligonucleotide is displaced by the refolding of a dsDNA fork. In this second situation, the fork is expelling the oligonucleotide thus reducing significantly its residence time. To account for our data in these two situations, we have constructed a mathematical model, based on the known nearest neighbor dinucleotide free energies, and provided a good estimate of the residence times of different oligonucleotides (DNA, RNA, LNA) with various length in different experimental conditions (force, temperature, buffer conditions, presence of mismatches, etc.). This study provides a foundation for the dynamics of oligonucleotide displacement, a process of importance in a number of biological and bioengineering contexts.

## INTRODUCTION

DNA hybridization is pervasive. It mainly happens under two general schemes. The first scheme involves oligonucleotides freely binding/unbinding on a ssDNA template. The other scheme involves competition between oligonucleotides on a DNA template. Both schemes play significant roles in fundamental biology. The free-binding configuration is known to be the foundation of base-pairing recognition, and the competition configuration plays a role in various biological conditions. During replication arrest (Fig.S1A), an oligo-hybrid can be displaced by the receding fork creating a structure known as a chicken foot that must be processed separately before replication can restart. Similarly during transcription (Fig.S1B), the competition between the DNA fork ahead of the transcription bubble and the DNA/RNA hybrid behind it may result in transcriptional pausing^1^. The competition between a fork and an adjacent hybrid has also been used in many practical applications, such as Hybridization Chain Reaction (HCR)^2^, DNA walker^3^, DNA Origami^4,5^, DNA sequencing^6^, and a genomic DNA editing technology^7^, known as Clustered Regularly Interspaced Short Palindromic Repeats (CRISPR) (Fig.S1C). CRISPR appears to rely on the opening of the DNA helix and the displacement of the forks by a complementary guide RNA oligonucleotide. In that instance, the efficiency and specificity of recognition and editing depends on the existence of mismatches between the guide RNA and the target DNA strands that might favour the displacement of the hybrid by the DNA fork(s). The competition and replacement of proteins bound on DNA is a similar process, which has been recently investigated in single molecule experiments^8,9^. It is thus important to characterize both hybridization configurations. To simplify the description, from now on, we refer to the free-binding situation as the ‘dissociation’ scheme and the competition situation as the ‘displacement’ scheme.

Previous studies have elucidated many aspects of the ‘dissociation’ scheme. By using Nearest Neighbour free energies^10–12^, one can accurately predict many equilibrium thermodynamics parameters, such as the melting temperature T_m_ and affinity constant K_d_. However, in many biological situations, the important parameter determining the occurrence of a reaction is the lifetime of a hybridized state but not its K_d_. Characterizing kinetic parameters, such as the hybridization lifetime τ_off_ or the binding time τ_on_, requires capturing the dynamic process of hybridization. Although fluorescent-based techniques (such as FRET^13^) have been used to investigate the kinetic parameters, due to the complexity of the experimental design, the data throughput is relatively low and the fluorescent labeling may affect the hybridization dynamics. In parallel, the kinetic parameters of the ‘displacement’ scheme remain uninvestigated, due technical difficulties in capturing real-time oligonucleotide competition.

In this paper, we propose a label-free platform to quantify the kinetic parameters for both schemes. Specifically, we used a magnetic trap to pull on a DNA hairpin. At a large enough tension, the hairpin is unzipped^6,14^ and the resulting ssDNA strand can hybridize with oligonucleotides in solution. As the tension on the molecule is reduced, the hairpin refolds but the DNA fork may be blocked transiently by the presence of these oligo-hybrids in the refolding pathway. With an oligonucleotide complementary to the hairpin stem, we characterized the ‘displacement’ kinetics by studying the blockage times during refolding, namely the displacement times, τ_disp_, by the receding fork of blocking oligo-hybrids as a function of force, temperature, length and type of oligonucleotide (e.g., RNA, or Locked Nucleic Acid (LNA), mismatches, etc.). Similarly, by using an oligonucleotide complementary to the hairpin apex, we characterized the ‘dissociation’ kinetics by studying the time lag required for hairpin nucleation and refolding upon tension release.

We found that the ‘dissociation’ kinetics is consistent with the standard oligonucleotide hybridization estimates, while the measured ‘displacement’ times are much shorter than the ‘dissociation’ times due to competition from the adjacent DNA from the hairpin. Additionally, at low forces, we observed that the existence of complementary sequences flanking the oligonucleotide allows for the possibility of encirclement at low forces thus reducing the blockage lifetime for hairpin nucleation and refolding.

We propose a model valid for the description of both ‘displacement’ and ‘dissociation’ schemes. By using the known free energies of hybridization between (complementary or non-complementary) bases, our model fits all the experimental data with a single adjustable parameter (the typical base pair unbinding time that sets the timescale). It is simple to exclude the encircling mechanism in this model thus providing a tool to predict the simple oligonucleotide dissociation. We also performed a stringent test of the model, sorting out the importance of various physical parameters, such as salt concentrations, by using an extra controlled variable (the tension on the fork). Together, this theoretical framework provides for new insights into this pervasive biological process.

## RESULTS

### A single molecule assay to study DNA fork blocking

We first focus on the ‘displacement’ scheme. Our assay is based on the mechanical unzipping of a DNA hairpin with a magnetic trap set-up (Fig.1A). Briefly, a DNA hairpin is used to tether a micron-size magnetic bead to a glass surface. Small permanent magnets are used to pull on the tethered bead and subsequently, above a threshold force, unzip the hairpin. This set-up allows both to control the applied force (via the distance of the magnets to the sample) and to measure the extension of the DNA molecule (by analyzing the image of the tethered bead) (1).

**Figure 1.**
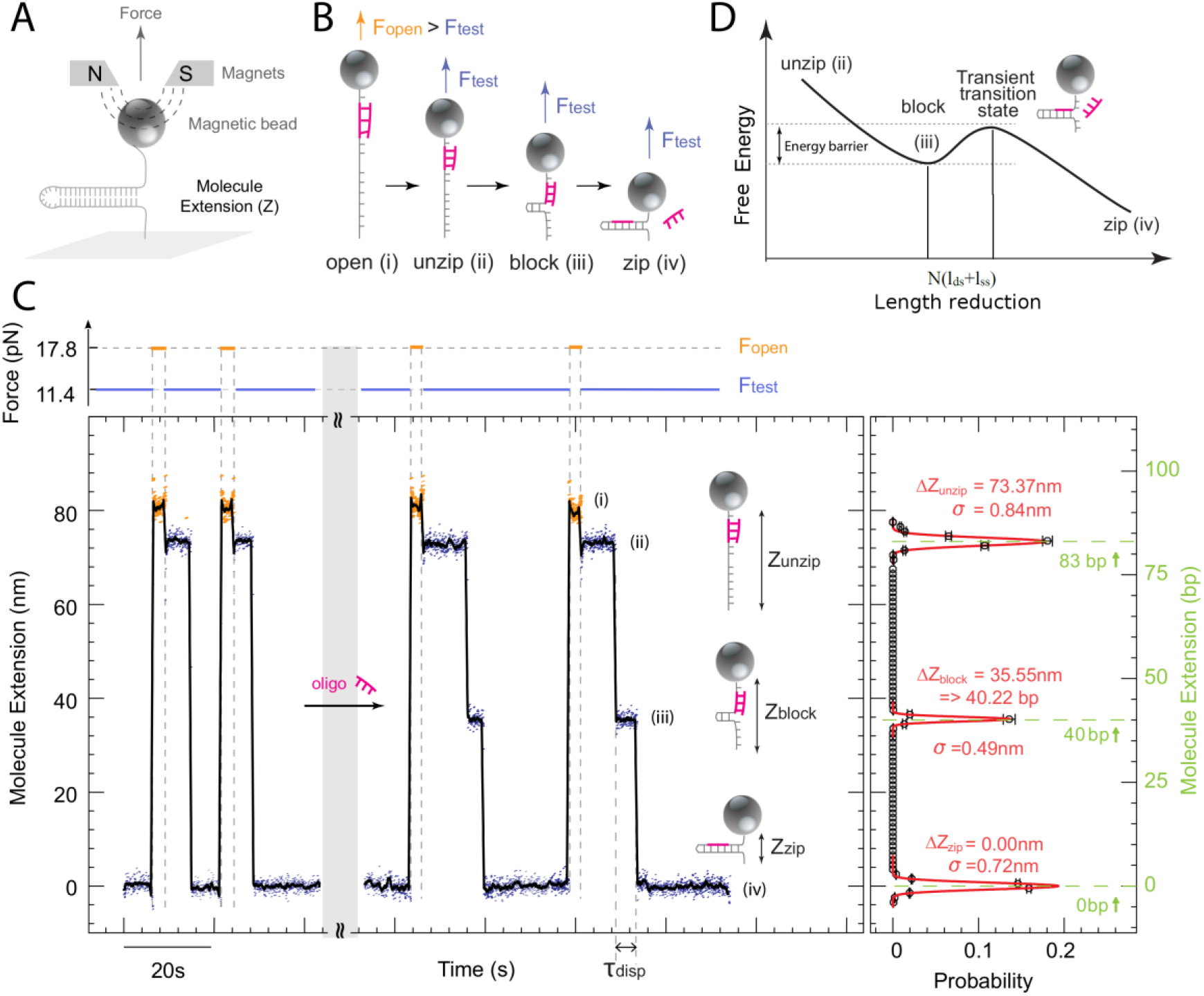
Detection of oligonucleotide-induced blockages during re-hybridization. (A) Schematic representation of the experimental set-up. (B) Schemes of the experimental process with the energy landscape. Five different extension levels are expected: (i) Z_open_ the fully unzipped hairpin at a force F_open_ large enough (>15pN) to open it, (ii) Z_unzip_, the transient extension of the unzipped hairpin at a force F_test_ too small (<12 pN) to prevent its re-hybridizaton, (iii) the partially rezipped hairpin blocked by an oligonucleotide at Z_block_, the transition state (iv) and (v) the extension Zzip of a fully folded hairpin. (C) Left panel: Experimental traces of an 83 bps hairpin recorded at F_open_ = 17.8 pN (orange) and F_test_ = 11.4 pN (blue; see force trace at the top), marked corresponding as in (B). The black curve corresponds to a 1 s average of the raw data. Right panel: Histogram of blockages. The black curve represents the histogram of the number of blockages per cycle at a given extension of the hairpin upon re-hybridization at F_test:_ ΔZ = Z_block_ -Z_zip_ in nm on the left scale and base pairs on the right scale, obtained from a single hairpin. Gaussian fits to the data are shown in red. The variance of these fits (σ ∼ 1 nm) defines the resolution of the apparatus. The roadblocks Z_block_ is observed at the expected positions (green dashed line). Notice that reducing the force from Fopen to Ftest results in a slight change in the extension of the ssDNA, due to its elastic properties (Fig 1C and Fig.S2)

In the fork-blocking assay, the DNA hairpin is first unzipped by applying a high force F_open_ (>15pN). Reducing this force to F_test_ (< 12 pN) allows for a quick refolding of the hairpins^15^.

Unzipping/re-zipping cycles can be repeated on the same molecule in the presence of a complementary oligonucleotide in solution. Hybridization events lead to transient blockage at F_test_, (see pause in the extension signal, Z_block_, at level (iii) in Fig.1B and 1C). Z_block_ reflects the actual position of the hybridized oligonucleotide (oligo-hybrid) on the DNA hairpin (conversion in bp is obtained using ssDNA elasticity as in^16–18^). This opens interesting perspectives for DNA identification and sequencing^6^. This configuration is similar to a stalled replication fork.

### Measuring the average displacement time <τ_disp_> in the fork-blocking assay

The oligonucleotide displacement time τ_disp_ is deduced from the statistics and duration of blockages. As shown in Fig. 2 (insert), these blockages display a single exponential distribution for τ_disp_ with a characteristic mean blockage time <τ_disp_>.

**Figure 2.**
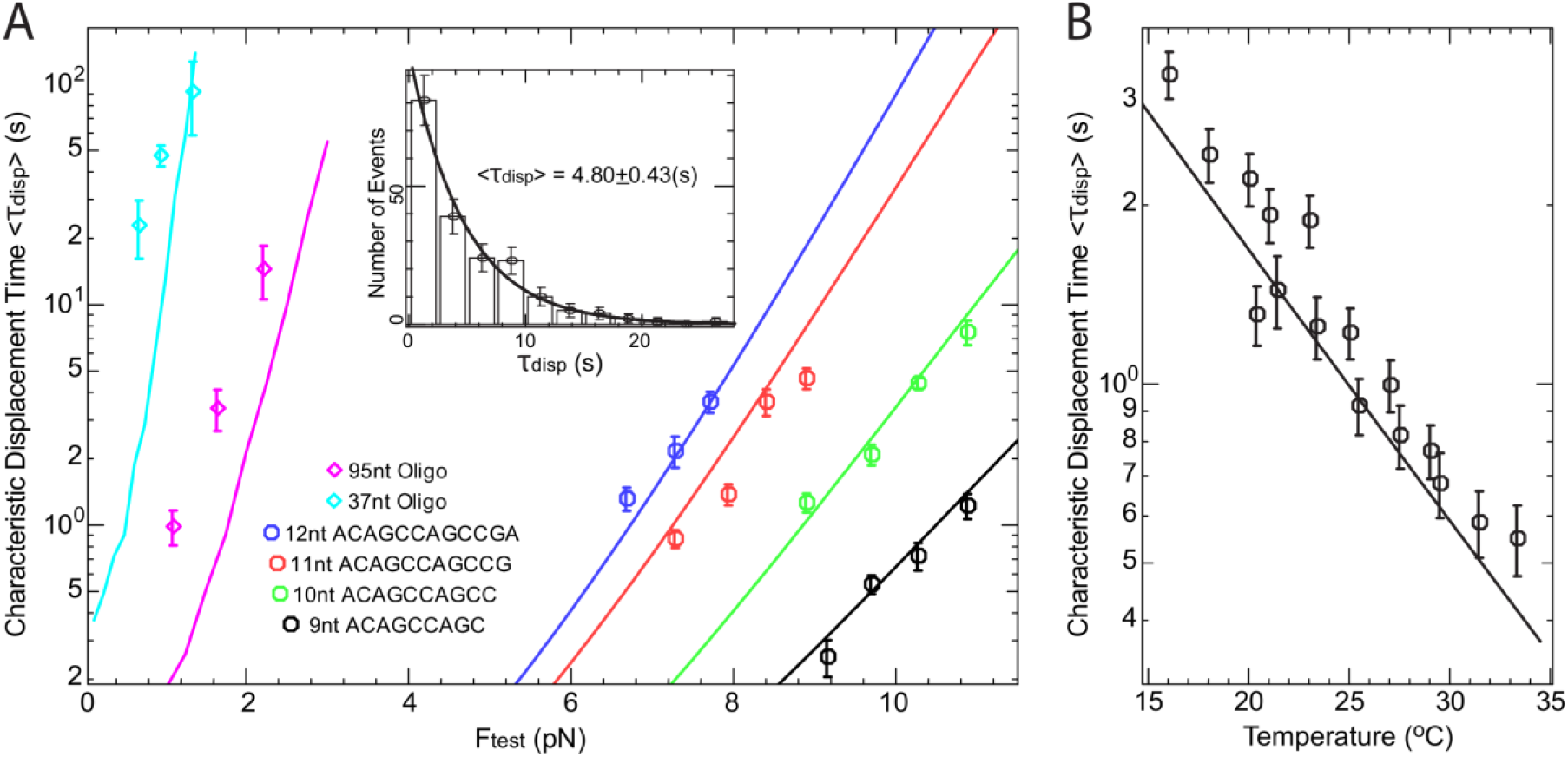
(A) The displacement time versus oligonucleotide length in the fork blocking assay. (Insert) The histogram of the blocking time τdisp displays a single exponential distribution with characteristic displacement time <τdisp>. τdisp are obtained from ∼200 fold/unfold hairpin cycles on the same molecule. The line is the prediction from a model of strand displacement (see Eq. 1). Points show the experimental evolution of **<**τ_disp_**>** versus F_test_ for oligonucleotides of length N (9 ≤ N ≤ 12). In a simple picture, **<**τ_disp_**>** varies exponentially with F_test_. As a result, for a given hairpin, **<**τ_disp_**>** can only be measured in a narrow force range. Adapting F_test_ provides a way to study the hybridization over a large range of lengths, here N varies from 9 to 95 nts. The colored continuous lines correspond to the predictions of the model described in the text. ((B) Displacement time for the 10 nt oligonucleotide at a force F = 8.5 pN as a function of the temperature. Decreasing the temperature increases the oligonucleotide stability as expected from the model (continuous line).

### Effect of the force F_test_, the oligonucleotide length N and temperature T on <τ_disp_>

The displacement time <τ_disp_> increases exponentially with both F_test_ and N and decreases exponentially with T (see Fig.2), a behavior consistent with our intuition: increasing F_test_ reduces the stability of the hairpin and its propensity to displace the oligo-hybrid; increasing the number of bases N in the oligo-hybrid increases the number steps required to displace the oligonucleotide; finally, high temperatures destabilize the oligo-hybrid. The displacement time <τ_disp_> is almost independent of the salt concentration (see Supp. Fig S12).

### Effect of the nature of the oligonucleotide and the presence of hybrid nucleotides and mismatches on <τ_disp_>

Nucleotides such as RNA, methylated bases and LNA improve the stability of their hybrid with DNA^19,20^ and thus increase <τ_disp_> : it increases by ∼4 for a 11 nt RNA oligonucleotide (Fig.3A) when compared to the same DNA oligonucleotide; a single LNA base increases <τ_disp_> by ∼2, and three LNA bases by ∼10 (Fig.3B); a single methylated cytosine may increase <τ_disp_> by ∼ 1.3 (data not shown).

**Figure 3.**
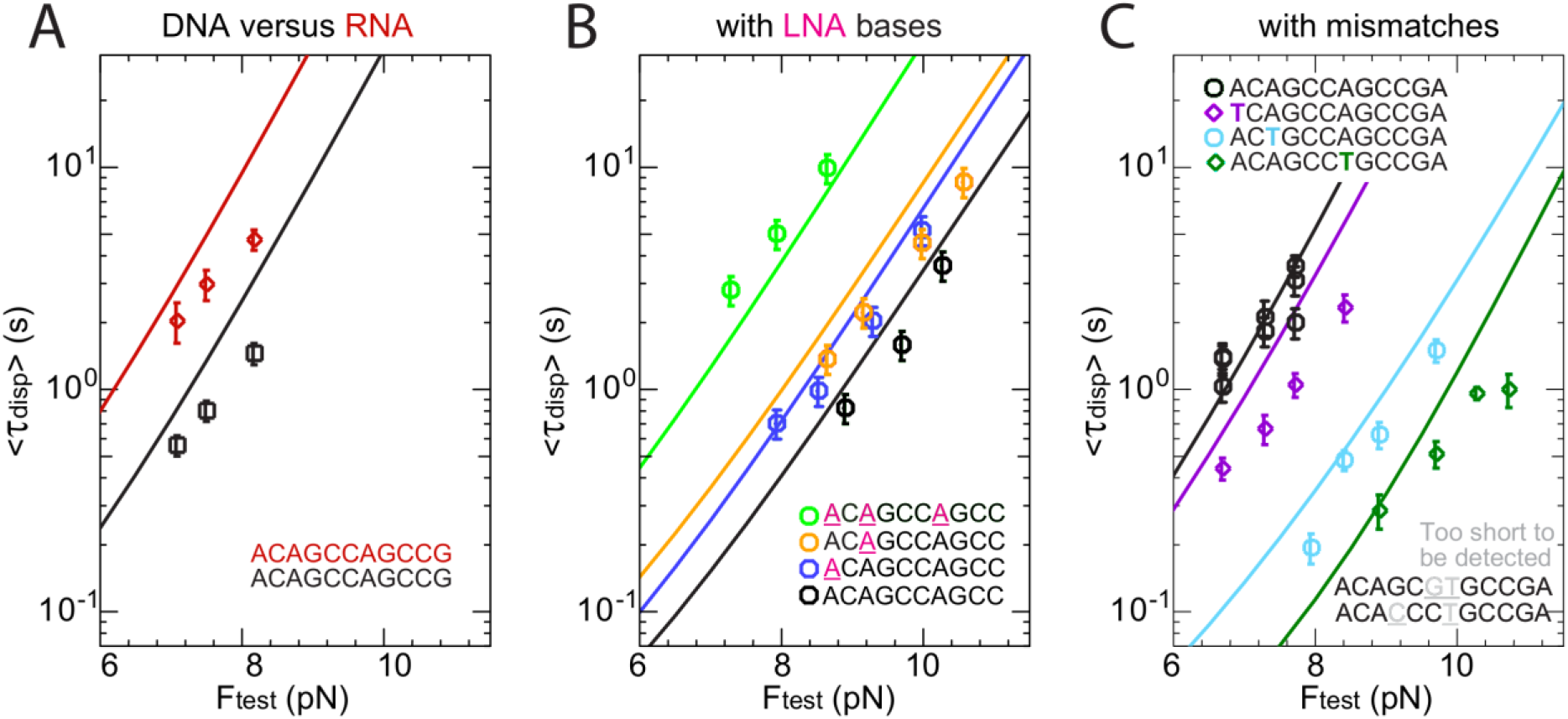
Influence of nucleotide types on the displacement time <τ_disp_>. (A-B) Increased binding stability induced by changing nucleotides from DNA to (A) RNA, (B) LNA base. The continuous lines are predictions based on the model described in Methods: (A) for 11nts RNA the model fits the data by considering an increase of stability of ΔΔG=1.1 kcal/mol for each the 11 nts DNA-RNA oligonucleotide, the prediction made using dinucleotide energy from^15^ overestimate the experimental results. (B) For 10nts oligonucleotides with 1 or 3 LNA bases. The model fits the data the following sequence dependent LNA-DNA increases in stability ΔΔG(AC) =-0.5 (Kcal/mol), ΔΔG(CA)=-0.1 (Kcal/mol), ΔΔG(AG)=-0.45 (Kcal/mol), the first and underlined base in the motif is a LNA (all full lines have been divided by 1.5 so that the pure DNA case fits properly). (C) Evolution of <τ_disp_> versus mismatch position in an oligonucleotide having 12 nts. The blue points correspond to the original oligonucleotide without mismatches, and the other colors correspond to an oligonucleotide with a mismatch at the underlined position. The model fits the data by considering, in agreement with^34^ a pairing parameter due to mismatches caused by the substitution of an A with a T in the oligonucleotide ΔG(TC/TG)=0.304 kcal/mol, ΔG(TG/TC)=-0.289 kcal/mol DG(CT/GT)=-0.289 kcal/mol. In agreement with the model the displacement time of the oligonucleotides ACAGCGTCCCGA and ACACCCTGCCGA with two mismatches are too short to be detected.

In contrast, mismatches reduce <τ_disp_> in a position-dependent manner. A mismatch at the first base reduces <τ_disp_> by a factor ∼5, and by a larger factor when in the middle of the oligo-hybrid. With two mismatches in the middle of a 12 nt oligonucleotide, blockage is no longer detectable within the resolution of our apparatus (i.e., <τ_disp_> < 100 msec).

Interestingly while the position of destabilizing mismatches strongly affects <τ_disp_>, stabilizing hybrid base pair (DNA-RNA, DNA-LNA, DNA-methylated DNA) is mostly position independent.

### The loop blocking assay: oligonucleotide binding to the hairpin apex

We now look into the ‘dissociation’ scheme that we call the loop blocking assay where the oligonucleotide being at the hairpin apex is not subject to the displacement by the fork (see Fig.4A). The oligo-hybrid inhibits refolding of the hairpin by sequestering its nucleation region. When the oligonucleotide spontaneously unbinds (*dissociation pathway*) after Fig.4A, the hairpin refolds (leading to a simple measure of τ_off_). At low forces an alternative *encircling pathway* appears, with the hairpin nucleating and refolding in the back of the blocking oligonucleotide, Fig.4A.

**Figure 4.**
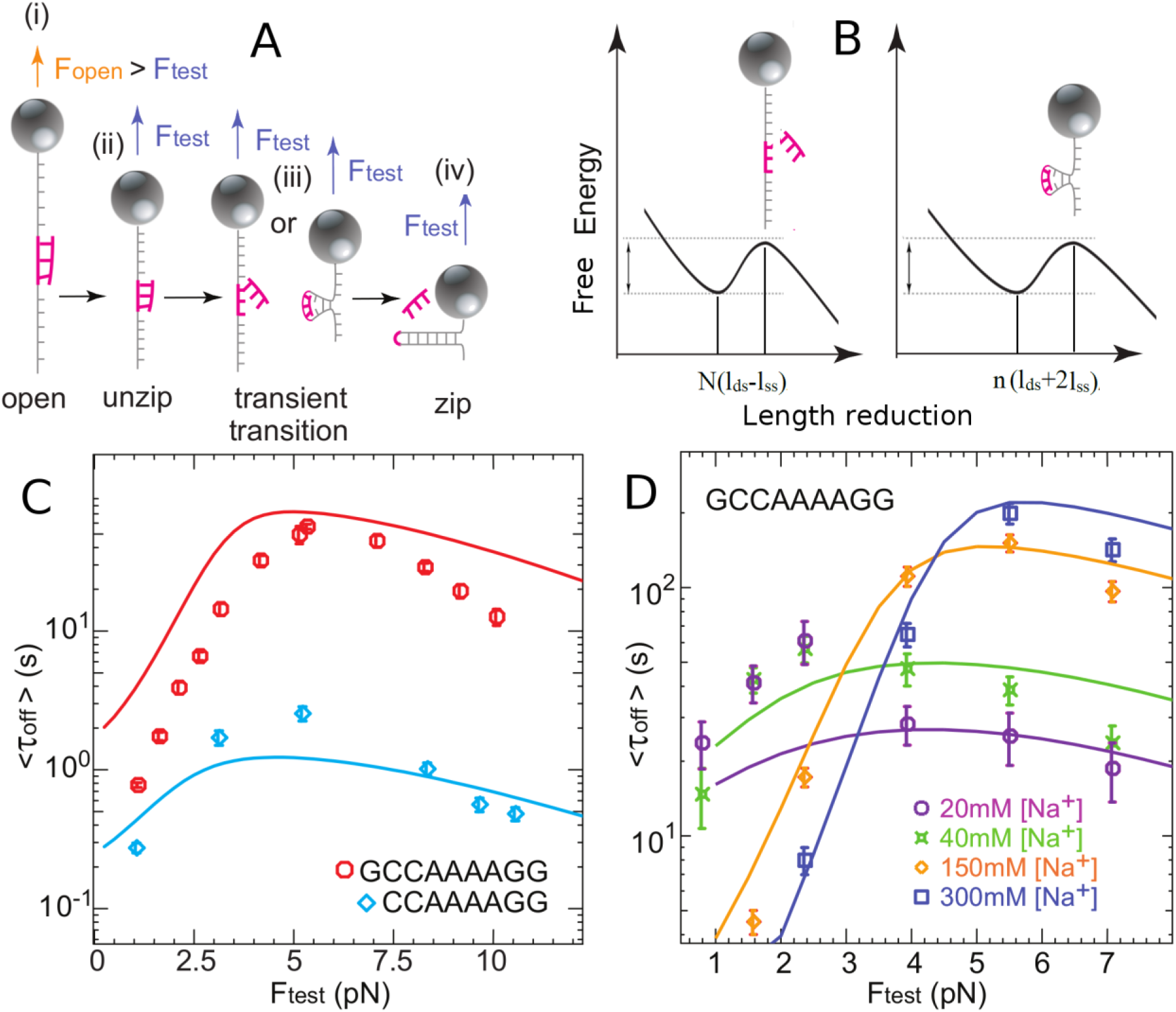
The oligonucleotide blocking time τ_off_ in the loop blocking assay versus F_test_. (A) Schemes of the experimental process (i) the hairpin is open allowing the oligonucleotide hybridization at the apex of the hairpin. When this hybridization occurs, it transiently prevents the hairpin refolding (ii) for a time τ_off_. Two mechanisms are possible for the oligonucleotide release: at low force (iii) the hairpin refolds encircling the oligonucleotide; at high force (iv) spontaneous detachment. The final state (v) corresponds to the hairpin refolded. (B) <τ_off_> versus F_test_ for an 8 nt (black) and 9 nt (blue) oligonucleotide blocking in the hairpin loop in Passivation Buffer (T = 25°C with [NaCl]=150 mM). Dotted lines are obtained by a polynomial fit of the data points. Full lines are predictions using the model in the main text without fork pressure. The energy barrier determining <τ_off_> is dominated by the pairing energies and depends on the force through the change of extension between ssDNA and dsDNA.

Due to the increase in <τ_off_> compared with <τ_disp_> and the range of timescales that can be explored (typically between 0.1 and 300 s), the length of the oligonucleotides studied is quite narrow: 8 or 9 nucleotides. At 5-6 pN, a 9 nt oligonucleotide displays a means dissociation time <τ_off_> twenty times longer than an 8 nt oligonucleotide, Fig.4C. We have studied the variation of <τ_off_> with F_test_ in various salts and temperature conditions. <τ_off_> increases by a factor ∼2 for a temperature decrease of only 3°C (see oligonucleotide GCCAAAAGG at 150 mM NaCl in Fig.4C at 25°C and Fig.4D at 22°C). Similarly <τ_off_> depends much more than <τ_disp_> on salt concentrations: at high forces (F>5 pN) the *dissociation pathway* leads to an increase of <τ_off_> by almost an order of magnitude between 20 and 300 mM NaCl. While at low forces the *encircling pathway* leads to a decrease of <τ_off_> by almost an order of magnitude between 20 and 300 mM NaCl.

### A model and a script predicting oligonucleotide displacement and dissociation

We modeled the displacement of an oligonucleotide by a closing dsDNA fork, in the blocking fork assay (“fork” mode), and its dissociation (“oligo” mode) or encirclement in the blocking loop assay (“apex” mode). In the three cases, the process is described by the base by base opening/closing dynamics driven by thermal fluctuations^21,22^. Opening/closing probabilities depend on elastic and base pairing energies (using the MeltingTemp.py script from BioPhyton 1.79) as described in Fig. 5. Such a coarse-grained model is either studied by Monte-Carlo simulations and or by a stochastic Transition Matrix approach.

**Figure 5:**
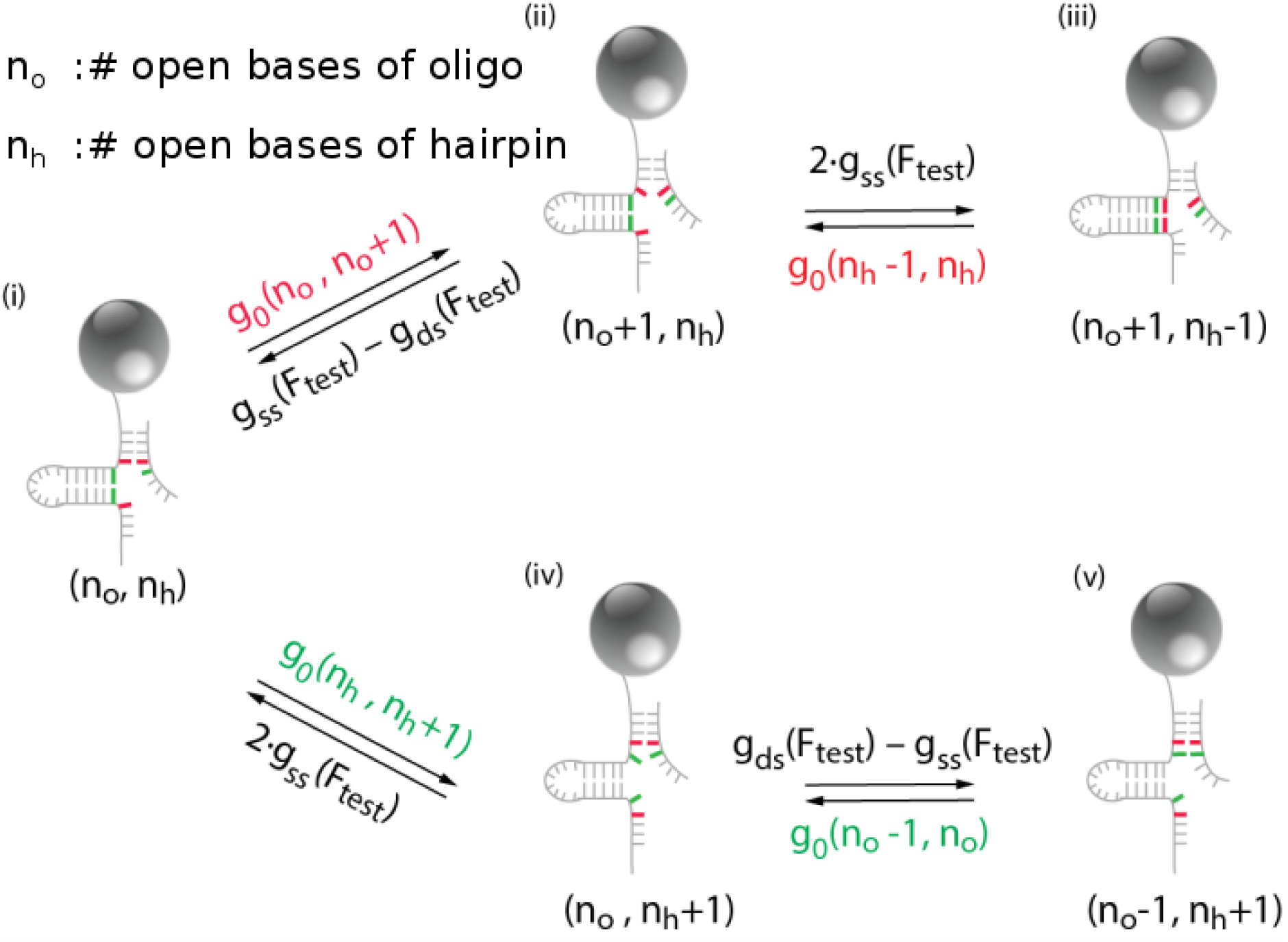
Some transition elements and how to escape from a (i) closing-blocked condition. Forward step: (ii) opening of the oligonucleotide followed by (iii) closing of the hairpin fork. Backward step: (iv) opening of the hairpin fork followed by (v) closing of the oligonucleotide. Note that the opening of a base pair of the oligonucleotide from (ii) and of the hairpin from (iv) are also possible but the direct closing of the oligonucleotide from (iii) and of the hairpin fork from (v) is not possible. gss (F_test_), and gds(F_test_) are respectively the single-strand and double-strand elastic energies at the pulling force F_test_. g_o_(n,n-1) are the zero-force base pairing energies which depend on the sequence (red or green bases) and which include base-stacking effects. The opening and closing rates in the transition matrix are proportional to the exponent of minus the energy costs associated with the transitions indicated by the arrows.

We provide a Python program corresponding to this last case which predicts <τ_disp_> or <τ_off_> for any sequence (with and without mismatches for DNA/DNA) and various salt conditions, forces, temperatures, and nucleotide types in the three modes (provide that the mismatch energy is known in BioPython).

**The displacement** (“fork” mode) is described by a scheme involving three fronts: both ends of the oligonucleotide experience peeling fluctuations, the hairpin fork being the third front. In practice, the oligonucleotide bases are most of the time fully hybridized, but fluctuations can drive the transient opening of one base. When an oligonucleotide base transiently opens on the side of the closing fork, the latter can advance, preventing further closing of the oligonucleotide (Figure 5). The oligonucleotide displacement is mostly achieved by the fork, while unpeeling at the opposite oligonucleotide end detaches one or two bases at most in the complete process (see Monte Carlo simulations in Supplementary materials Fig. S6).

Each time the fork advances by one base, there is no net loss in pairing energy because the base pairing is exchanged between the oligonucleotide and the hairpin complementary strand. Thanks to this replacement mechanism the displacement process is very fast compared to spontaneous dissociation. It has indeed an overall energetic cost only due to the shortening of the pulled hairpin strand under the force F_test_. Such activation barrier (see Fig.1D explains via an Arrhenius law the observed exponential distribution of the displacement times <τ_disp_ > (see Fig.2A (Inset)), its increase with the force F_test_ and the oligonucleotide length, and the possibility to displace very long oligonucleotide by lowering F_test_ as observed in the experiments (Fig 2A). Mismatches and base variants can also easily be taken into account by modifying the pairing energy in the replacement process (Fig 3).

**The spontaneous dissociation** pathway (“oligo” and “apex” mode) in the loop blocking assay is described by an unpeeling process (“oligo” mode), which can be combined with encirclement (‘apex’ mode). Unpeeling engages the two ends of the oligonucleotide^23^ At each step, the position of the first or the last hybridized base of the oligonucleotide has the possibility to unpeel or re-zip by one base. Owing to the difference in energy, the probability for a re-hybridization (which only costs the difference in the stretching energy at F_test_ between double and single strand) is much higher than for unpeeling (which cost the energy of the base pairing). The unpeeling fronts stay most of the time close to the first and the last bases. But on some rare occasions a series of unpeeling events occur that lead to complete unbinding of the oligonucleotide.

As shown in Fig.4B&4C the large activation barrier due to unpairing (dissociation), describes the general behaviour of the experimental results at large forces and the great dependence of the displacement time on oligonucleotide length and sequence. To describe the rapid decrease of <τ_off_> observed at low forces we have added in our model an encircling pathway, which accounts for the possibility that a hairpin stem forms downstream of the oligonucleotide with a cost depending on the loop formation energy, resulting in the hairpin encircling the hybrid and displacing it.

As shown in Fig. 4 a model combining unpeeling and encircling (“apex” mode) in the transition matrix fits the data well. There is a crossover force between the two processes at about 5 pN depending on a model for the elementary encircling rate (see methods and Supplementary material).

### Comparison of the experimental results with our model prediction

The coarse-grained description introduced here reproduces our experimental data with only few adjustable parameters: t_0_ an elementary time for each step (which sets the timescale for displacement), λ a salt correction factor, β and t_u_ (t_u_ = t_0_ 0.31^n^) the salt correction and elementary rate in encircling (see Supp. Mat. Section x). Note that while t_0_ and λ are involved in all modes, t_u_ and β are only required in the encircling mechanism occurring in the apex mode. We have used the pairing-energy parameters, from the literature^10–12,24–26^, and suggested some corrections for the salt dependence and hybrids base pairs interactions (DNA-RNA, DNA-LNA). We have set the parameters defining the elasticity of dsDNA and ssDNA from the literature and our measurements^16^.

While Fig.2-4 presents a comparison with the “fork” and “apex” modes, Fig. 6 provides a comparison with experiments done by other groups^13,27,28^ looking at simple oligonucleotide hybridization (“oligo” mode).

**Figure 6.**
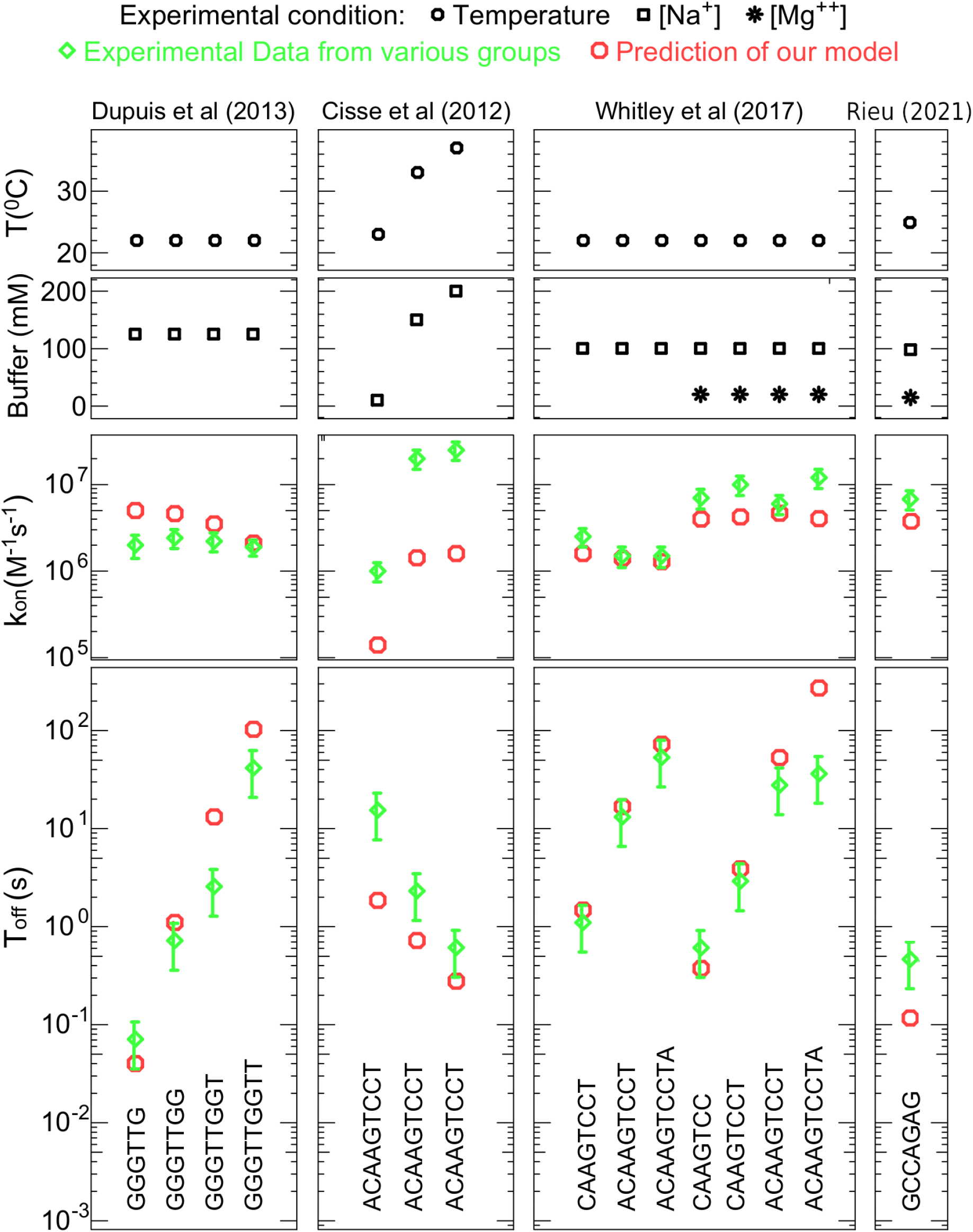
Comparison of simple oligonucleotide binding kinetics with experimental measurements done by various groups, Dupuis et al 2013^27^, Cisse et al^13^, Chemla et al^28^ and Rieu et al^38^. The quality of the agreement depends mostly with duration of τ_off_, all these measurements (excepted Rieu) are made using fluorescence which is limited by photobleaching.

As can be seen on Figures 2 and 3, the agreement is good for <τ_disp_> with a t_0_ = 2.5 µs. The agreement is less good when using RNA or LNA published ΔGs^15,20^. For RNA, the prediction of τ_disp_ is too high and too low for LNA. In the comparison of an oligonucleotide containing either all bases in RNA or in DNA, the predicted ΔΔG^25^ reaches 1.58 k_B_T for 11 nt while our best fit is obtained for 1 k_B_T (0.6 kcal/mol). These energy differences lie within the errors of the nearest neighbour’s parameters. The conclusion is the same for LNA. In fork mode <τ_disp_ > does not depend significantly on the salt concentration but the binding probability increases proportionally with it. Our model predicts a mild increase of 50% of the displacement time when salt increases from 50 mM to 300 mM. We understand this effect since each base pair opening of the oligonucleotide is replaced by the same base pair closing in the hairpin.

In the “apex” mode, t_0_ = 1 µs, this replacement disappears and <τ_off_ > increases with the salt concentration (Figure 4B), as already reported in 18. Our model fits well this dependency if we decrease by a factor λ ∼ 0.5 the salt correction normally applied to ΔS when calculating the oligonucleotide melting temperature^10,29^. This reduction is probably due to the difference in Manning condensation between dsDNA and ssDNA^30^ (see Supp. Data).

This salt effect is reinforced in the encircling pathway as the two ssDNA strands need to hybridize. We need to enhance the salt correction on ΔS by a factor β ∼2 to correctly fit the experimental data.

So far we have discussed the k_off_ = 1/T_off_ of an oligonucleotide, but we can also access the k_on_: using the free energy of the base pairs and the normal salt correction (λ = 1) we compute the ΔG of the oligonucleotide and its K_d_ = exp(ΔG/RT). Then we access to k_on_ using k_on_ = k_off_/K_d_. Zhang et al^5^ have measured the K_on_ of a large set of 36 nt oligonucleotides and proposed a model predicting this K_on_. Compared with Fig. 6.

## DISCUSSION

Our assay allows measuring the kinetic of unbinding and replacement of an unlabeled oligonucleotide, in contrasts with other approaches which use a fluorescently labeled oligonucleotide and thus are prone to the limiting observation time caused by photobleaching and to changes of binding affinity due to the interactions of fluorescent dyes with nucleic acids^31,32^.

The oligonucleotide displacement described here is very similar to the discovery of the replacement process of proteins bound to DNA by proteins in solutions^9^, acting much faster than spontaneous dissociation^22,33^. Here the replacement of the oligonucleotide is achieved by the DNA closing fork; there is then no dependence on the concentration of competing molecules in <τ_off_> but the set-up allows to use of the testing force as an essential control parameter.

The displacement model introduced here is an extension of the model for unzipping of DNA under force^33,34^ but includes the competition of two unzipping processes on the same DNA. This model describes replacement at a base-pair scale which is more detailed than the strand displacement^35^, the multi-strand simulator^36^, or the single pathway models^37^, but less detailed and computationally much less costly than molecular dynamics simulations^37^ (see Supplementary material for more details).

Our model provides a reasonable agreement with our experiments taking into account that these times vary exponentially with the physical parameters involved.

When using an oligonucleotide complementary to a sequence in the hairpin apex, we have been able to investigate the unbinding time of the oligo-hybrid, a situation which has been investigated quite extensively using other means (mostly using fluorescently labelled oligonucleotides^13,27,28,38^, though rarely with unlabeled ones^28,38^). We have used the same model to analyse these published experimental data. The agreement between the model predictions and the published data is reasonable as most of our predictions are within a factor ∼4 of the experimental values. The most severe disagreements occur for long τ_off_ (> 100 s) where the experimentally reported value is shorter than our prediction. That discrepancy could be attributed to photobleaching which affects the estimate of long τ_off_. The experimental data might also be affected by interactions between the fluorescent dyes and the DNA which may affect the hybridization free energies as reported in^32^. Overall, our model provides a means to estimate the hybridization time of oligonucleotides to their template with and without mismatches and it provides a solid foundation for the understanding of oligo-hybrid displacement by a receding fork, a situation of biological importance.

We have also observed that at low forces, nucleation of a hairpin in the back of a blocking oligonucleotide could account quantitatively for our observations. Such processes could play an important role in other situations where formation of a DNA bubble around an enzyme could mask its action, as in the unwinding of a DNA fork by a helicase^39^ or the formation of a R-loop.

The sensitivity of <τ_off_>, the nature and position of mismatches have also been recently reported in a pure hybridization assay^6^. When using the present assay to probe a DNA sequence^6^ such sensitivity could be used to detect SNP ‘Single Nucleotide Polymorphism’^6^.

Our work provides a tool to predict the dissociation time τ_off_ of an oligonucleotide which should be useful for many applications. The present accuracy is reasonable but there is room for improvements by adjusting the various parameters for which we have proposed starting values, this could be done by using more elaborate ssDNA elasticity model^17,18^ and by using systematic label-free measurement as proposed in^38^.

## CONCLUSION

The novel assay proposed here provides insight on the displacement of DNA/DNA hybrids by a complementary strand, a situation encountered both in-vivo such as in transcription pausing and replication arrest and in vitro during DNA-origami formation^4^ and the study of DNA walkers and CRISPR^7^. Using this approach, we have already successfully achieved DNA sequencing and identification at the single molecule level^6^. We provide here a model and a software allowing predicting the oligonucleotide displacement time and thus to optimize probes for these applications. We also demonstrate its validity by measuring the oligonucleotide unbinding kinetics. As a further application, the novel assay combined with the replacement model can be used to better determine base-pair parameters and end effects in pairing energies.

## MATERIALS AND METHODS

### Magnetic Tweezers

Experiments were done on a commercial PicoTwist magnetic tweezers (www.picotwist.com) with single DNA hairpins tethered between a coverslip via a digoxigenin-anti-digoxigenin bond and a magnetic bead via a biotin-streptavidin bond. DNA molecules were mixed with streptavidin magnetic beads (Dynabeads^@^ MyOne™ C1, Invitrogen). Then, we gently flowed the mixture into a 5mm x 40 mm x 40 um pre-coated anti-digoxigenin chamber. After 5-15mins incubation, we rinsed thoroughly the chamber to remove unattached beads/samples. We changed the chamber environment to defined buffer condition with proper oligonucleotides respectively. The applied force (F_test_, F_open_) on the hairpin is controlled by varying the distance of a pair of permanent magnets from the sample. The position and extension of the hairpins are monitored through a 30 Hz video-camera (CM-140 GE Jai) and tracked through a self-written program. More details on magnetic tweezers (including the flow cell fabrication, sample preparation and molecule manipulation as well as measurement) can be found in^6^.

### Single molecule assay

Due to the high degree of parallelism of magnetic tweezers, we can simultaneously apply the same force on many single molecules. For every experiment in this paper, we always repeatedly zip-unzip molecules >100 times, deduced the needed value (e.g. <τ_disp_>, P_block_, etc.) for each molecule and calculated the variation (i.e. error bars in Fig.2-4,6) among different molecules. Unless marked otherwise, all the experiments were performed at room temperature under T4 buffer condition: 0.2% BSA (Bovine Serum Albumin), 25 mM Tris−Ac, 150 mM KOAc, 10 mM MgOAc_2_, filtered by 0.22 μm filter. The oligonucleotides are ordered from Eurogentec and Integrated DNA Technologies. The detailed protocol to synthesize 1.2 kbps hairpin is described in^40^. The preparation method for 83 bps hairpin can be found in^6^.

### Oligonucleotides sequences

The sequences of small (<12 nucleotides (nt)) are given in the Figures. The sequence of the 37 nt oligonucleotide is: 5’-GGGTGTTTGATTGATTTGATTCCTTGGATGTGCGAAG-3’ The sequence of the 95 nt oligonucleotide is :5’TTTGGCTAAATATCCTAATGTTAAAG TGTATGATAAGCCTACTACAGTAGATTTTGACGGGTGTTTGATTGATTTGATTC CTTGGATGTG CGAAG-3’

### Elasticity model for ssDNA, dsDNA and pairing energy parameters

For the stretching energy g_ss_(f) of a ssDNA base at the force f we use the freely jointed-chain model with monomeric length b_0_=1.828 Å and effective stretching length d=0.605 Å (S1) For the stretching energy g_ds_(f) of a dsDNA base pair at the force f we use the Odijk formula^41^ for the Worm-Like chain model with stretching corrections (S2) with monomeric length h=3.4 Å, and the persistence length is ξ=50 nm.

### Nearest Neighbour parameters

The pairing parameters g_o_(n,n+1) depend on the base pair and the following one due to stacking effect, and can be decomposed in the enthalpic and entropic contribution : g_o_(n,n+1)=ΔH(a_n_,a_n+1_)+TΔS (a_n_,a_n+1_,ic). Where a_n_ is the nucleotide in position n on the oligonucleotide and ic is the ionic conditions. We have taken the ΔH and ΔS parameters from Santa Lucia, see Supplementary Tables. We have not used any initialization cost but an end destabilization obtained by halving the pairing energy of the last two attached base pairs of the oligonucleotide. Note that we estimate such positions to be at the antepenultimate and penultimate position for the fork blocking assay and to be in the middle one for the case of the loop blocking assay for the two oligonucleotides under study which have a symmetric sequence.

### Dissociation time of the oligonucleotide in the fork-blocking assay

The model, as detailed in Supplementary Material, is based on a stochastic transition matrix built from the probabilities to open or close a base pair of the oligonucleotide (on one of the two sides) or of the hairpin, until the oligonucleotide expulsion.

As described in Fig 5 The probability to open base pairs n_o_ of the oligonucleotide depends on its pairing and stacking energies, g_o_(n_o_,n_o_+1), (^29,42^ and Supplementary Tables S3 and S4) r_oO_(no)= 1/to exp(-g_o_(n_o_,n_o_+1)), where the probability to close a base pair depends only on the force via the single strand and double strand stretching energies, g_ss_(F_test_) and g_ds_(F_test_)): r_cO_= 1/to exp(-g_ss_(F_test_)-g_ds_(F_test_) The probability to open the base pair n_h_ for the hairpin is r_o_ ^h^(n_h_)= 1/to exp(-go(n_h_,n_h_+1)), while the probability to close a base pair is r_ch_= 1/to exp(-2g_ss_(f)). The transition matrix is of dimension growing as N^3^, where N is the number of oligonucleotide base pairs, due to the 3 possible opening-closing fronts, it is zero for opening-closing transitions on non-adjacent base pairs, it has heterogeneous entries due to the sequence dependence of the pairing energies, moreover, reflect the fact that closing of a base is not possible if it is already paired with the competing random walk (See Supplementary).

In the standard replacement process, for a complementary DNA oligonucleotide, there is no overall lost in pairing energy. The total cost to replace, by the closing fork, N nucleotides occupied by the oligonucleotide is:

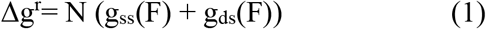

we then expect for the Arrhenius law the exponential decay observed exponentially:

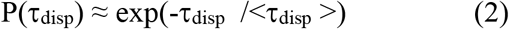

where ln <τ_disp_ > ∝ ΔG_disp_ /k_B_T. and d(ln <τ_disp_ >) /dF ∝ N(l_ds_+l_ss_)/k_B_T, l_ds_(F) and l_ss_ are the double-strand and single strand extension per nucleotide at the force F (Supplementary). Such barrier (1) shown in Fig. 2D explains the observed dependence of <τ_disp_ > on the oligonucleotide length and the possibility to replace very long oligonucleotides by lowering the force Ftest. Such description is not sufficient to precisely calculate <τ_disp_> and to study its dependence on temperature, ionic conditions, mismatches or hybrid bases or RNA in the oligonucleotides. The solid lines for <τ_disp_> in Fig. 2 &3 are indeed obtained by numerical diagonalization (**9 ≤ N ≤ 12**) or Monte-Carlo simulations (N=37 and 45) of the transition matrix defined by the above rates.

### Dissociation time of the oligonucleotide in the loop-blocking assay

To calculate the dissociation time of the oligonucleotide in the loop-blocking force we model the unpeeling of the oligonucleotide by a stochastic step-by-step unpeeling from both the 5’ and 3’ ends of the oligonucleotide, with opening rate r_oo_(i)= 1/to exp(−go(i,i+1)) and closing rate r_co_= 1/to exp(−g_ds_(f) + g_ss_(f)).

The Arrhenius energy barrier for this process is

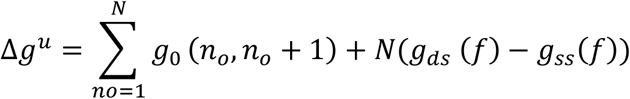

The first term, which does not appear in the replacement energy barrier (eq 1), corresponds to the unpairing energy of the oligonucleotide at zero force and is the sum of the N nearest-neighbour pairing parameters g_0_(i,i+1)^20,42^. This contribution is large and explains the large stability of the oligonucleotide: one additional pair in the 9 bp oligonucleotide with respect to the 8 bp oligonucleotide, *g*_*0*_ *(G,C)* ∼*3 k*_*B*_*T*, gives the measured ratio between the unbinding times e^3^∼20. With the same reasoning, the expected time for a 7 bp oligonucleotide will be in the range of hundreds of milliseconds, in the limit of our detection range. The second contribution to the unpeeling energetic barrier in (eq 3) derives from the non-zero test force, and is the difference between the stretching energy of the double strand and the single-strand DNA. Using the Arrhenius description (like in eq 2) we obtain d(ln<τ_off_ >) /dF ∝ N(l_ds_-l_ss_)/ k_B_T

Such a force behaviour explains the presence of a maximum in τ_off_ around 5 pN in Fig. 4C (where l_ds_ = l_ss_). At f >12 pN *g*_*ds*_ (*f*) > *g*_*ss*_(*f*) and the unpeeling is facilitated by the stretching force, but it is still an activated process, due to the base-pairing term. Only at very large forces of several tens of pN (40 to 60 depending on sequences) the activation barrier disappears^23^. As detailed in methods, similarly to the replacement, we have precisely calculated the τ_off_ of the oligonucleotide using a stochastic base by base unpeeling process occurring from both 3’ and 5’ ends^23^.

To model the energetic barrier for the encirclement Δg^c^ we suppose that the minimal encircling loop with *n* hybrid hairpin-oligonucleotide base pairs in the middle is a triangle with *n* single-strand base pairs on each side of the double strand (see the transition state b) in Fig. 4) and we compute the mechanical energy required to shorten the extension of the molecule by such triangle:

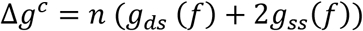

and, using (2) d(ln<τ_off_ >) /dF ∝ n (l_ds_+2 l_ss_)/ k_B_T.

## ACKNOWLEDGMENTS

We thank B. Ducos, C. Andre, for useful comments on the manuscript and R. Monasson for helpful discussions on the modelling. This work was supported by ERC grant Magreps 267 862 (to V.C.), by the Fondation Pierre-Gilles de Gennes, program FPGG032 (to V.C.), by the Centre National de la Recherche Scientifique.

