## Supplementary Materials for "Displacement and dissociation of oligonucleotides during DNA hairpin closure under strain"

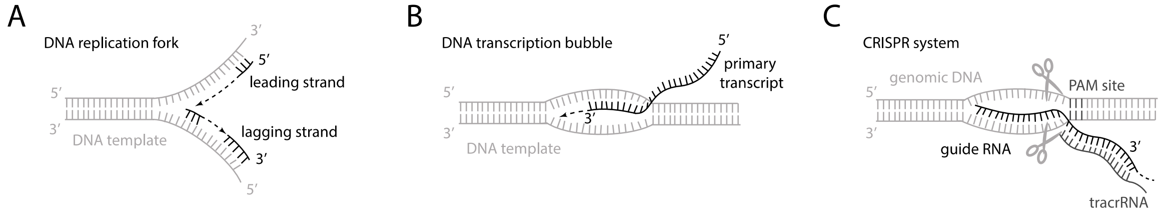


**Fig. S1.** The scheme of three biological examples involving the competition between a DNA fork and its adjacent hybrid(s).

#### Elements of Unzipping-Unpeeling models

The elastic energy of a single DNA strand (ssDNA) of N bases at force f is, with a freely jointed chain model:

$$G_{ss}(N,u) = N g_{ss}(u) = N\, b_o log\left[\frac{\sinh(u)}{u}\right] + \frac{0.5\, f^2}{s\, k_B T} \qquad \text{(S1)}$$

where s=216 pN , u = d f /$k_B$ T. We use the parameters for the ionic conditions of 10 mM NaCl: d = 0.542 nm, $b_o$ =2.14 nm.

The freely-jointed-chain free energy (S1) as a function of the force and the corresponding force-extension curve are plotted in Fig.S2. The experimental force-extension curve shown in Fig.S2 is very well reproduced by the above model, except at large forces (f >15 pN), out of the range of interest, where the elasticity of the backbone has to be introduced in (S1) for a correct modeling.

The free energy of the double-strand DNA (dsDNA) at fixed force is described by the Odijk formula for the Worm-Like chain model with stretching corrections

$$G_{ds}(N,f) = N\, g_{ds}(f) = N\, h\left[\frac{f}{k_B T} - \sqrt{\frac{f}{A\, k_B T}} + \frac{0.5\, f^2}{s' k_B T}\right] \quad (S2)$$

where h = 0.34 nm and the persistence length is A=50 nm and s'=1230 pN.

The pairing parameters $g_o(n,n+1)$ depend on the sequence and buffer conditions (ionic condition, presence or not of magnesium, temperature) are taken from the server DNA Fold (1,2) .

The extension of a ssDNA base, $l_{ss}(f)$, and of a dsDNA base pair, $l_{ds}(f)$, at force f are obtained by deriving the free energy (S1,S2) with respect to the force.

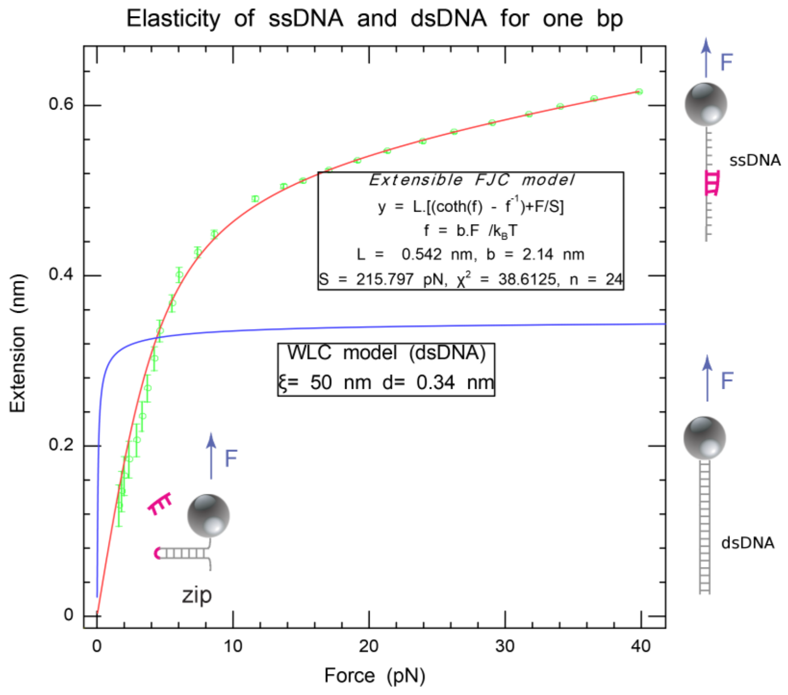


**Fig.S2:** Experimental force-extension curve for ssDNA nucleotides and dsDNA base pairs and fit. The green curve of ssDNA is obtained by a protocol explained in Gutierrez et al [8]

.

### Transition Matrix for the Fork blocking assay

The model used to calculate the replacement time of the oligonucleotide with the hairpin fork is sketched in Fig.6 of main Text and briefly described in the method section, here we give a more detailed description.

The model is a zipping model: at each time step the hairpin fork and the oligonucleotide can stochastically open or close by one base on the pulled DNA hairpin. The closing of the hairpin and of the oligonucleotide are in competition because both the sequences are complementary to the one strand of the hairpin which is pulled, therefore closing of a base pair of oligonucleotides/or hairpin fork is not possible when its complementary base on the pulled strand is already paired with the hairpin fork/ or oligonucleotide respectively.

Starting from the configurations with the opening front in position the oligonucleotide open or close a base with transition rates: $r_o^o(i) = e^{-go(i-1,i)}$ and $r_c^o(f) = e^{-gds\,(f)+gss\,(f)}$ (3). When a base pair opens the pairing energy is indeed lost, while when it closes the stretching energy difference between the final double strand configuration and the initial single strand configuration, at force f, is lost. The hairpin unzips and rezips with transition rates, $r_o^h(i) = e^{-go(i-1,i)}$, $r_c^h(f) = e^{-2gss(f)}$. Note that because of the closed hairpin is perpendicular to the force when a base pair is closed the stretching free energy of the two open bases is lost.

We describe each configuration by 3 indexes ($i_{5'}$,$i_f$,$i_{3'}$) representing the position of the opening fronts (on the 5'-3' oligonucleotide): $i_{5'}$ is the position of the 5' front, $i_f$ the position of the fork and $i_{3'}$ the position of the opposite 3' front . The replacement process is a stochastic dynamics described by the equation:

$$\frac{\partial P(i5',iF,i3')}{dt} = M\,P(i5',iF,i3') \quad (S3)$$

Where $P(i_{5'},i_F,i_{3'})$ is the probability that the 3 fronts are in position $(i5', iF, i3')$ .

An example of transition matrix for a 3-base oligonucleotide is given in Table S1. More generally non-zero elements of the transition matrix are: $M\,[(i_{5'},i_F-1,i_{3'})\ ,\ (i_{5'},i_F,i_{3'})]=r_o^h(i_F)$, $M[(i_{5'},i_F+1,i_{3'}),\ (i_{5'},i_F,i_{3'})]=r_c^h(f)$, $M[[(i_{5'}+1,i_F,i_{3'})\ ,\ (i_{5'},i_F,i_{3'})]]=r_o^o(i_f)$, $M[[(i_{5'}-1,i_F,i_{3'})\ ,\ (i_{5'},i_F,i_{3'})]]=r_c^o(f)$, $M[[(i_{5'},i_F,i_{3'}-1)\ ,\ (i_{5'},i_F,i_{3'})]]= r_o^o(i_{3'})$, $M[[(i_{5'},i_F,i_{3'}+1)\ ,\ (i_{5'},i_F,i_{3'})]]= r_c^o(f)$. The Diagonal element $M[[(i_{5'},i_f,i_{3'})\ ,\ (i_{5'},i_f,i_{3'})]]$ is minus the overall rate of leaving the configuration $(i_{5'},i_F,\ i_{3'})$. Notice that once the oligonucleotide is detached it cannot reattach (and the hairpin can rapidly close). Transition to the oligonucleotide deshibrydation is therefore escaping transition, only contributing to the diagonal terms. All opening transition from the position in which the 5' and 5' front are the same position $i_{3'}=i_{5'}$ are escaping transitions.

An important condition is that to close a base pair of oligonucleotides or of the hairpin fork the complementary base not to be already paired. This can be summarized by the condition that the hairpin fork cannot be at the same position of the 5' or 3' oligonucleotide front. All such configurations are forbidden and this reduces the number of elements of the matrix.

The closing rate only depends on the force and is different for the hairpin fork and the oligonucleotides unpeeling fronts $r_c^O= 1/to\ \exp(-\ g_{ss}(F_{test})-\ g_{ds}(F_{test})$, $r_c^h= 1/to\ \exp(-\ 2g_{ss}(f)\ )$.

Where to is the elementary time for opening/closing fluctuations and is fitted to = α=2.5 μs

The opening rates are given by $r_o^o(i)= 1/to\ \exp(-g_o^o(i\text{-}1,i))$, $r_o^h(i)= 1/to\ \exp(-g_o^h(i\text{-}1,i))$. The nearest neighbor pairing parameters $g_o^o(i\text{-}1,i)$ and $g_o^h(i\text{-}1,i)$ are the same for standard DNA-DNA pairing, but are different for mismatches, LNA-DNA and RNA-DNA hybrids.

The pairing parameters are calculated for the specific oligonucleotide sequence, read in the standard 5',3' direction from the parameters given in Table S3 and S4 for standard DNA-DNA oligonucleotides or non-standard RNA-DNA,LNA-DNA or for mismatches. They have been derived from melting experiments, see Santa Lucia Hicks (2004) Annu. Rev. Biophys and available using BioPython software (https://biopython.org/) or IDT server .

We have taken the correction of AT base pairs terminal as in Santa Lucia Hicks (2004) Annu. Rev. Biophys

The border conditions, mimicking the presence of an opening fork which is not only extended on one base pair is that the pairing energies are divided by 2 when there are only 2 or less closed base pairs.

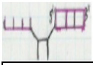

| | | | | | | | | | | |
|---|---|---|---|---|---|---|---|---|---|---|
| | 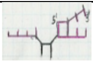 101 | 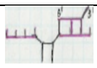 102 | 103 | 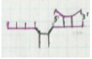 202 | 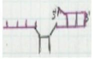 203 | 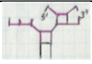 212 | 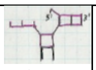 213 | 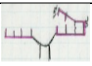 303 | 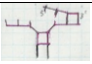 313 | 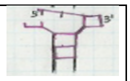 |

| | | | | | | | | | | |
|---|---|---|---|---|---|---|---|---|---|---|
| | | | | | | | | | | 323 |
| 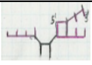<br>101 | $-r_c^o$<br>**$-2r_o^o(1)$** | $r_o^o(2)$ | | | | | | | | |
| 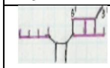<br>102 | $r_c^o$ | $-r_o^o(1)$<br>$-ro^o(2)$<br>$-r_c^o$ | $r_o^o(3)$ | $r_c^o$ | | | | | | |
| 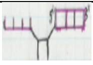<br>103 | | $r_c^o$ | $-r_o^o(1)$<br>$-r_o^o(3)$ | | $r_c^o$ | | | | | |
| 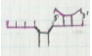<br>202 | | $r_o^o(1)$ | | $-2r_c^o$<br>**-**<br>**$2r_o^o(2)$**<br>$-r_c^h$ | $r_o^o(3)$ | $r_o^h(1)$ | | | | |
| 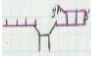<br>203 | | | $r_o^o(1)$ | $r_c^o$ | $-r_c^o$<br>$-r_o^o(2)$<br>$-r_o^o(3)$<br>$-r_c^h$ | $r_c^o$ | $r_o^h(1)$ | $r_c^o$ | | |
| 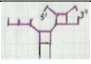<br>212 | | | | $r_c^h$ | | $-2r_c^o$<br>$-r_o^h(1)$<br>**-**<br>**$2r_o^o(2)$** | $ro^o(3)$ | | | |
| 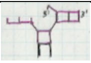<br>213 | | | | | $r_c^h$ | $r_c^o$ | $-r_o^o(2)$<br>$-r_o^o(3)$<br>$-r_o^h(1)$ | | $r_c^o$ | |
| 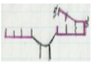<br>303 | | | | | $r_o^o(2)$ | | | $-r_c^o$<br>**$-2r_o^o(3)$**<br>$-r_c^h$ | $r_o^h(1)$ | |
| 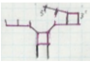<br>313 | | | | | | | $r_o^o(2)$ | $r_c^h$ | $-r_o^h(1)$<br>$-r_c^o$<br>**$-2r_o^o(3)$**<br>$-r_c^h$ | $r_o^h(2)$ |
| 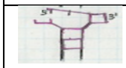<br>323 | | | | | | | | | $r_c^h$ | $-r_o^h(2)$<br>**$-r_o^o(3)$** |

**Table S1**: Transition Matrix for the Fork Blocking assay and for an oligonucleotide of 3 bases. The allowed configurations are sketched, and labeled by the 3 positions $i_{5'}, i_F, i_{3'}=1..3$, of the unpeeling front on the 5'direction, the hairpin fork and the unpeeling front on the 3' directions. The escaping transitions are in bold.

#### Transition Matrix for the Apex blocking assay

| | 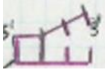<br>11 | 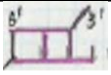<br>12 | 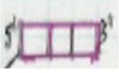<br>13 | 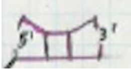<br>22 | 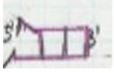<br>23 | 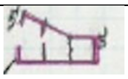<br>33 |
|---|---|---|---|---|---|---|
| 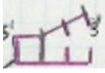 | **$-r_o^o(1)$-**<br>$r_c^l(f)$ | $r_o^o(2)$ | | | | |

| | | | | | | |
|---|---|---|---|---|---|---|
| 11 | | | | | | |
| 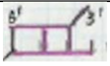 12 | $r_c^o(f)$ | $-r_o^o(1)-$ $r_c^o(f)-$ $r_o^o(2)$ | $r_o^o(3)$ | $r_c^l(f)$ | | |
| 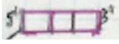 13 | | $r_c^o(f)$ | $-r_o^o(1)-$ $r_o^o(3)$ | | $r_c^o(f)$ | |
| 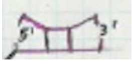 22 | | $r_o^o(1)$ | | **$-r^c(n=1,f)$** **$-r_o^o(2)$** $-2\ r_c^o(f)$ | $r_o^o(3)$ | |
| 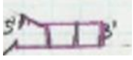 23 | | | $r_o^o(1)$ | $r_c^o(f)$ | $-r_o^o(3)$ $-r_c^o(f)$ $-r_o^o(2)$ | $r_c^o(f)$ |
| 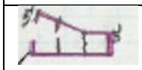 22 | | | | | $r_o^o(2)$ | **$-r_o^o(3)$** $-\ r_c^o(f)$ |

**Table S2**: Transition Matrix for the Loop Blocking assay and for an oligonucleotide of 3 bases. The allowed configurations are sketched, and labeled by the 2 positions $i_1$, $i_2$ = 1..3, of the unpeeling fronts on the 5' and 3' directions. The escaping transitions are in bold.

The Transition Matrix for the Loop Blocking assay has the same opening and closing transitions described above, and is shown for a oligonucleotide of 3 bases in TableS2. The opening rate for a fork in position i (indicated by the configuration index) is defined by $r_o^o(i)$= 1/to exp(-go(i-1,i)) and closing rate $r_c^o$= 1/to exp( $-g_{ds}(f) + g_{ss}(f)$).

The escaping transition due to encircling has a probability $r^c(n,f)$=1/tu exp($-\Delta g^c$) with energetic cost

$$\Delta g^c = n\ (g_{ds}\ (f) + 2g_{ss}(f) \quad \text{(S4)}$$

depending on the length n of the encircling loop (given by 2n simple strand bases and n double strand bases) and on the force F.

The elementary transition for the encirclement is slower than the elementary rate and it is exponentially decreasing with on the length of the encircling loop, we have fitted it to

$t_u$ = to $0.31^n$ .

#### Calculation of the displacement time of the oligonucleotide

To calculate the average displacement time we want to count the average time spent on all configurations with the oligonucleotide hybridized:

$$< \tau_{disp} >= \int_0^\infty dt\ p(\,oligo\ attached) = \int_0^\infty dt\ \ v_{hd} e^{M't}\ v'_o = -v_{hd} M'^{-1} v'_o \quad \text{(S4)}$$

where $v'_o$ is the initial condition *i.e.* the oligonucleotide is completely attached, while the hairpin fork is completely open: the only component of $v_o$ equal to one correspond to $i_{5'}$=1,$i_{3'}$=N-1, where N is the number of base pairs of the oligonucleotide. Note that $e^{M't}$ is the matrix exponential, and $M'^{-1}$is the inverse matrix. $v_{hd}$ is the vector which takes into account all the 'holding' configurations with at least 1 closed base pair for the oligonucleotide : all the components of $v_{hd}$ are one because we have removed the absorbing state of the detached oligonucleotide. We therefore compute the displacement time by (S4) from the numerical inversion of the matrix M and its scalar product with the holding and initial configuration.

#### Animation for a simple replacement process:

**Olidisp.svg**: is a Scalable Vector Graphic (SVG) animation describing the fork blocking assay. This animation runs in most web browsers just by opening this file with the browser (Firefox, IE, …), in some case your browser may ask you if you agree to run the script, just reply yes. The animation should look like:

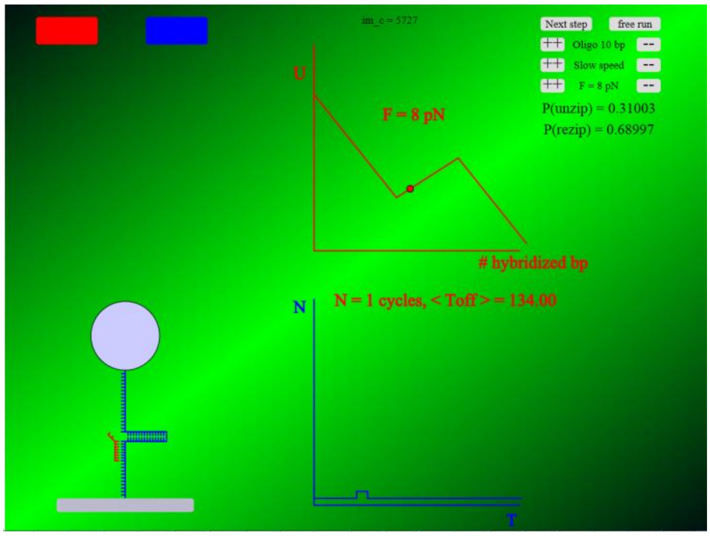


**Fig. S3** : The left part is a sketch of the assay and the state of the hairpin. In the top right, you will find buttons allowing modifying the oligonucleotide size, the force and the speed of the animation. The red curve illustrates the energy landscape and a point corresponding to the hairpin state. The blue graph displays a histogram of the $T_{disp}$, to observe this graph it is best to transiently specify the fastest animation rate and then reduce this rate. One then observes an exponential distribution as follows:

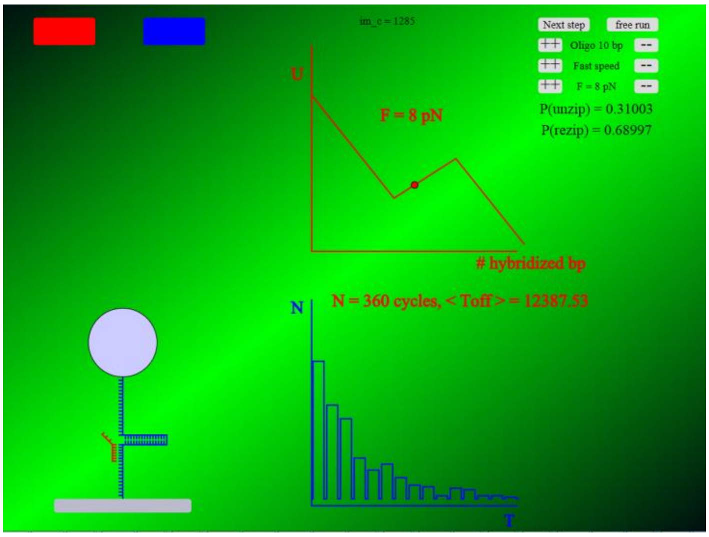


**Fig. S4** :This animation relies on a simplified version of the Monte Carlo simulation software (in SVG and javascript) with all bases being equivalent and no peeling of the oligonucleotide at its end opposed to the hairpin fork. The time is in arbitrary units in this animation.
Web animation illustration of the oligonucleotide blocking assay: this svg script should display in most web browsers (Firefox, Internet Explorer,…). This animation displays the assay in a semi-realistic manner: the DNA hairpin mimics the real one with dsDNA always having the same extension (which is realistic if F > 1 pN) and ssDNA which extension is force-dependent with a simple form. The magnets are not to scale nor their displacements. The user can modify the parameters by clicking on ++ and – buttons in the top right corner. He has the ability to modify the oligonucleotide length, the display speed and the force in the re-zipping phase. When modifying the force parameter, the unzipping probability Pu and re-zipping probability Pr are computed in a simplified way were all nucleotides are equivalent and assuming that Pu + Pr = 1. In the top right, the simplified potential energy graph of the system is displayed in red to help understand the phenomenon. In the bottom right, a cumulative histogram presents the distribution of blocking times versus this blocking time. This histogram should converge towards an exponential when the numbers of cycles N is large enough. If the user modifies either the oligonucleotide length or the force, the histogram is reinitialized. Selecting the display speed to the smallest rate allows to see all the steps in detail, the hairpin opening and closing is far too slow compared to the real experiment. Switching the display rate to tultra-fast, skips the hairpin detailed evolution and allows to have many cycles done and observe a meaning full histogram. Although the results of this animation follow the expected trends the values of the displacement time is not realistic.

### Monte Carlo simulations of the displacement kinetics:

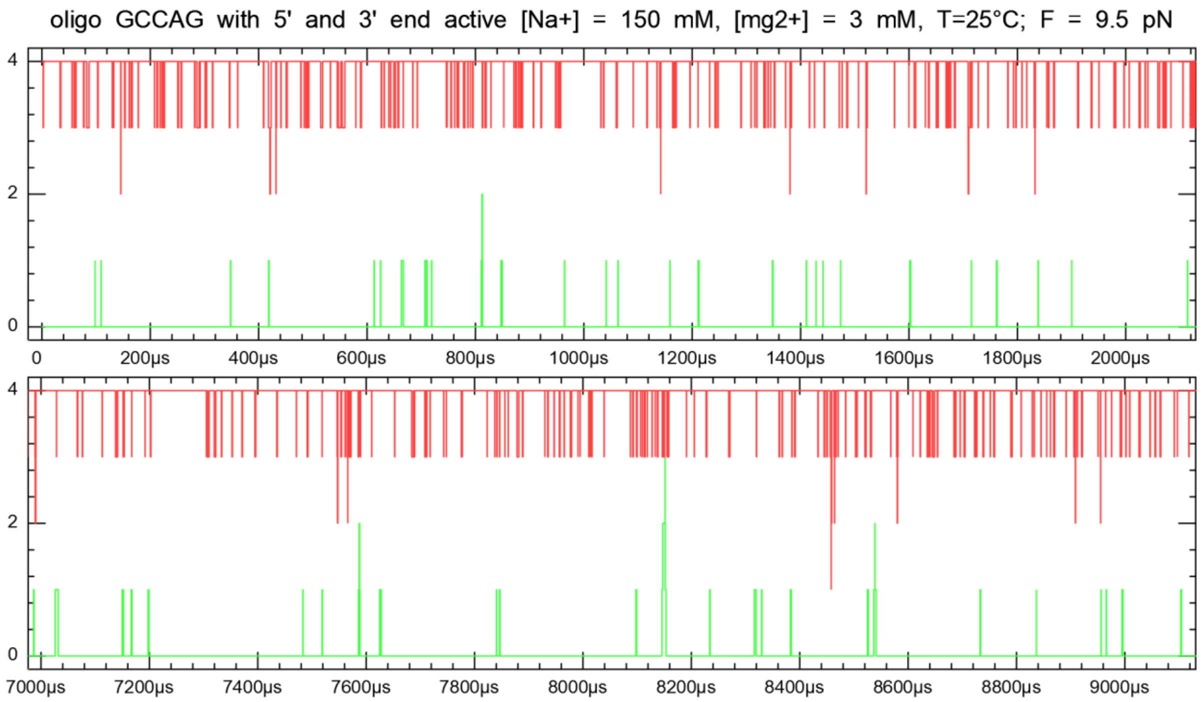


**Figure: S5** : Monte Carlo simulation of a 5 nts single oligonucleotide hybridized to its template under tension. Most of the time the oligonucleotide is fully hybridized, in frequent events the first or the last di-nucleotide opens transiently, in some rare occasion the next di-nucleotide also opens transiently, we thus have two fronts invading stochastically the oligonucleotide. To detach, the two fronts need to collide, at 7580 µs an aborted collision occurs. A more dramatic event occurs at 8150 µs, but the two fronts did not collide. The $\tau_{off}$ of this oligonucleotide equals 7660 µs. Notice that the 3' front corresponding to the AG di-nucleotide invades the oligonucleotide more frequently then the 5' front corresponding to the GC di-nucleotide.

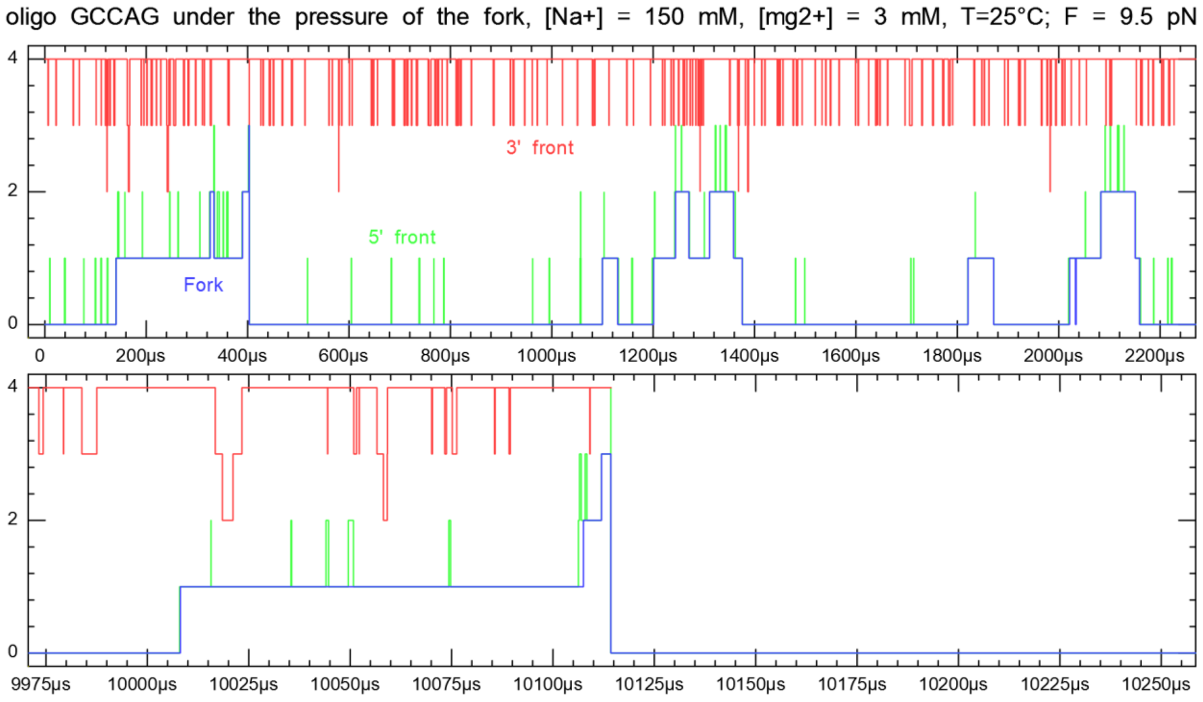


**Figure: S6**: Monte Carlo simulation of a 5 nts single oligonucleotide hybridized to its template under the pressure of the fork trying to close the hairpin. Compared to the previous case, the pressure of the fork breaks the symmetry.

The 5’ end of the oligonucleotide which is the more stable because GC rich, opens for a significant time stabilized by the fork. The oligonucleotide detachment occurs with the progression of the 5’ front reaching the 3’ end.

We have develop a Monte Carlo program reproducing the dynamics of the oligonucleotide with the same conditions as the Python program that is an oligonucleotide displaced by a fork under tension or just hybridized to a template under tension. We have use exactly the same parameters as those used in the Python program. The first thing that we have done it to check that both programs give the same results. With the Python program you get directly the $\tau_{disp}$ or $\tau_{off}$, with the Monte Carlo you can access to a description of the dynamics but each displacement or dissociation event is stochastic and leads to a random team with an exponential distribution with a mean time equal to the Python program result. Thus to accurately test the equality of the results between the two programs we need a substantial number of events since the accuracy on the mean time scales like the root square of the number of events. To reach a good accuracy we have used a GPU to run many Monte Carlo simulations in parallel (in Cuda). We choose the DNA oligo 5'-ACAGCCAGC-3' with 9 bases, T=25°C, F=10pN; Na+=100mM; Mg++=0mM; as a test. Our Python program leads to $\tau_{disp}$ = 0.611182 s. Our Monte Carlo program with 1638400 detachment events gives $\tau_{disp}$ = 0.6112 ± 0.0005 s, which is a good agreement. The distribution is indeed exponential (data not shown) but this is not true at a very short time. Indeed as the very large number of events demonstrates the oligonucleotide needs a minimal time to dehybridize which make sense as shown in the Figure S7 below for this 9 nts it takes at least ~350 µs.

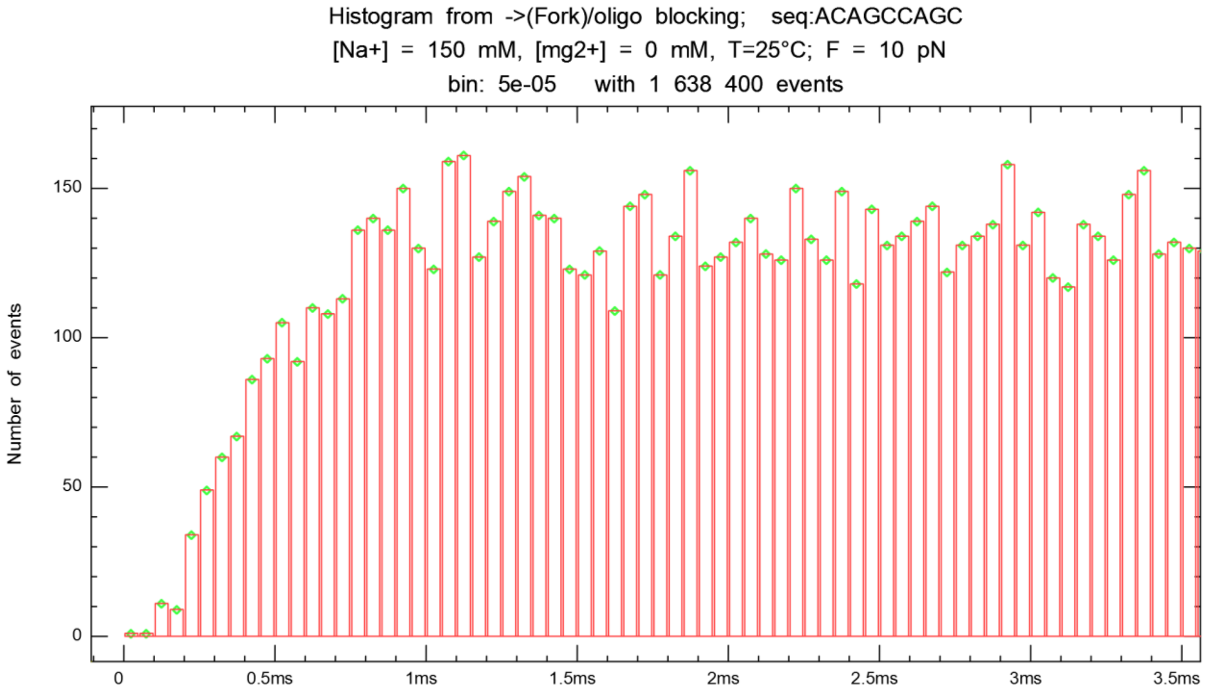


**Figure S7** : Histogram of the detachment times of the ACAGCCAGC oligo under the pressure of a fork under 10 pN unzipping force focused on very short events (at long time the distribution is exponential). With a very large number of events, we observe that the process requires a minimal time (350 µs) to occur.

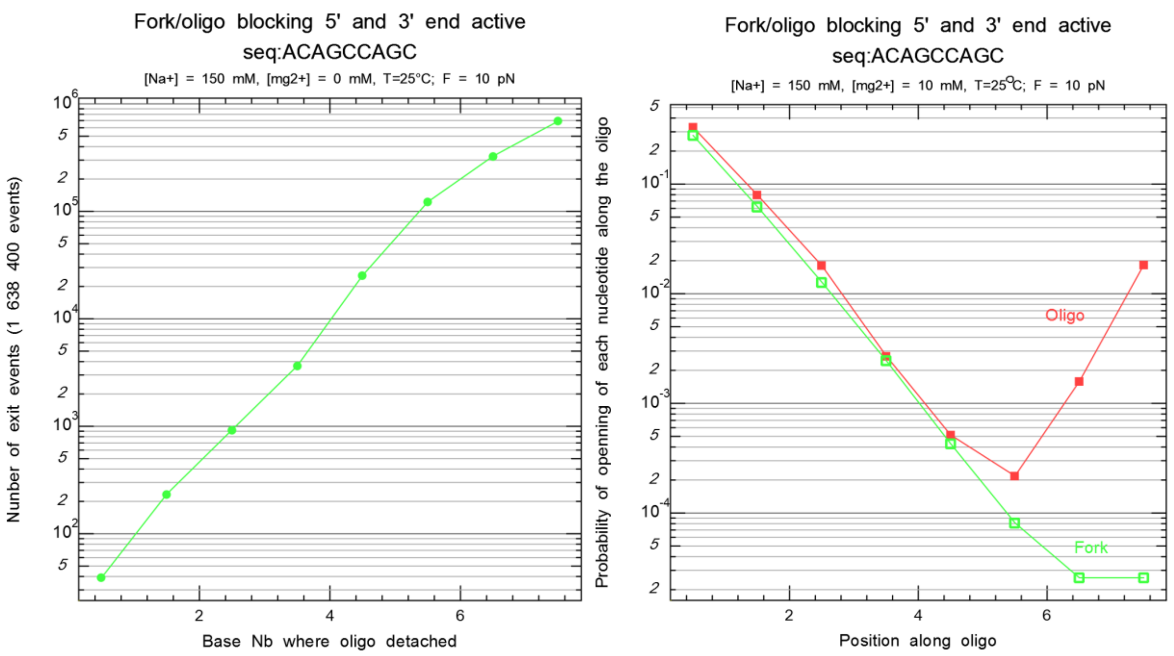


**Figure S8** : The Monte Carlo simulation of the displacement of an oligonucleotide by a fork shows that the fork is actively expelling the oligonucleotide these results contrasts with the dissociation of an oligonucleotide hybridized with a template under tension.

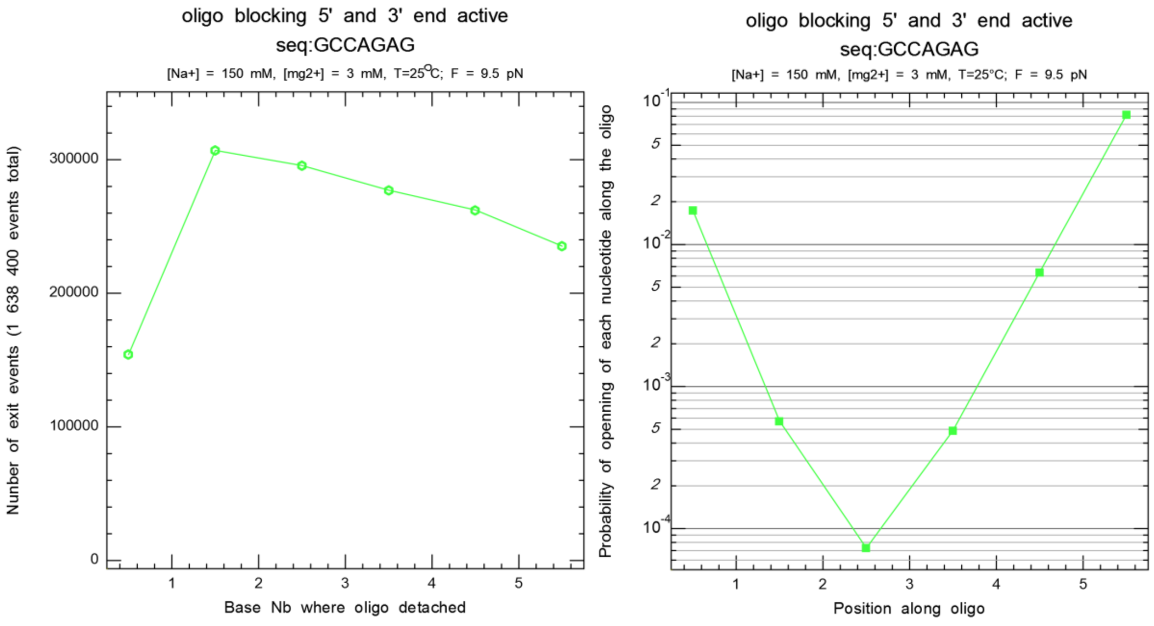


**Figure S9** : The Monte Carlo simulation of the dissociation of an oligonucleotide from its complementary strand under tension shows that the fork the oligonucleotide opens mostly at the ends and that the last hybridized base is not very specific.

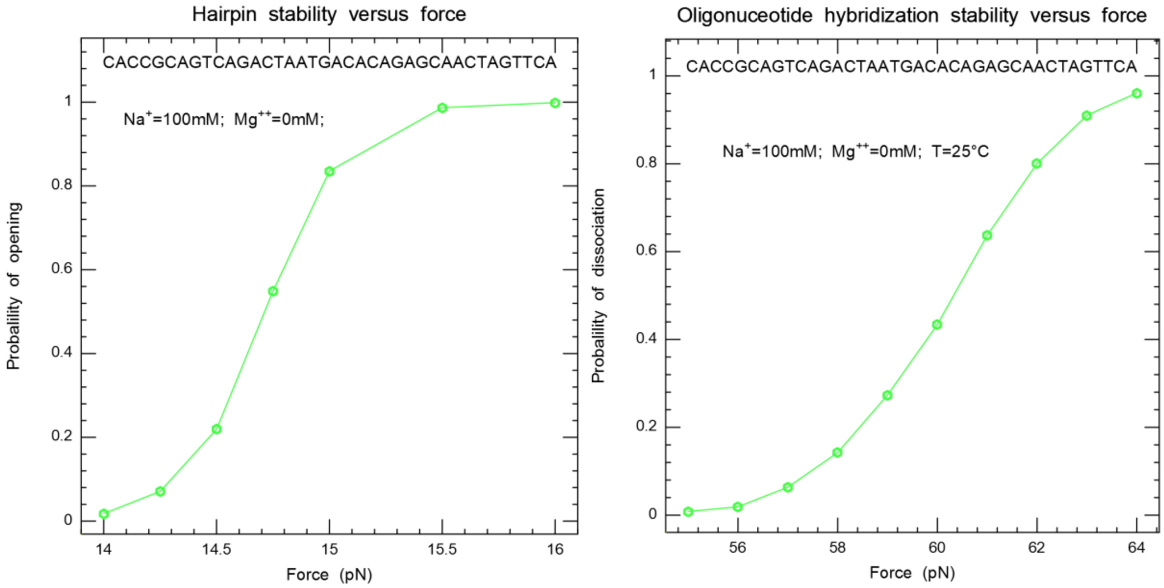


**Fig. S10**: Test of the integrity of the model: Using a Monte Carlo simulation program with exactly the same parameters as the Python program we develop, we test two situations which corresponds to know experimental results. We have applied the test to the same sequence of 38 nts part of the sequence chosen by Manosas et al xx which is known to be 50% AT rich and present a very flat energy landscape. In the first simulation, the initial condition corresponds to a half-open hairpin where the 19 nts on the 5' end are hybridized while a unzipping force is applied on the two strands. On the left panel, we present the opening probability versus force. We find that the hairpin opens at 14.7 pN which is in agreement with the known unzipping force. In the second panel, we challenge the hybridization stability of the same oligonucleotide partly hybridized with its complementary template which is under tension. This corresponds to the unpeeling transition which is known to occur experimentally around 60 pN. The initial condition of the Monte Carlo situation corresponds to have 19 nts of the middle part of the oligonucleotide hybridized with the template under tension while both ends of the oligonucleotide are not hybridized over ¼ of the oligonucleotide. The right panel presents the probability of dissociation of the oligonucleotide versus force, the transition occurs around 60.3 pN. Both results ensure that the elasticity model sused are reasonable.

A Monte Carlo simulation of the displacement kinetics is shown in Fig.S5 from the Eq. (S5) for different pulling forces, showing the attempts before the expulsion of the oligonucleotide and closing of the hairpin (see also the animation of the replacement process). The theoretical calculation of the replacement time is in agreement with the simulation.

We have used the Monte Carlo simulations to check the program based on the analytical calculation. Moreover for the long oligonucleotides with N=37 and N=95 nucleotides, it is computationally more convenient to calculate the replacement time by the Monte Carlo simulation described above.

#### Effect of sequences on the hybridization kinetics of DNA

As discussed in our model, the displacement time of the oligonucleotide does not depend exponentially on the total pairing energy of the sequence due to the replacement mechanism.

However the elementary fork displacement time of a base depends slightly on the sequence due to the prefactor $e^{g_0}$ to open a blocking closed base pairs before the displacement of the fork. As shown in Fig.S5 the dynamic zipping model reproduces the slight variations of the average displacement time with the sequence for an 11 bp oligonucleotide, these variations do not depend simply on the GC content of the sequence.

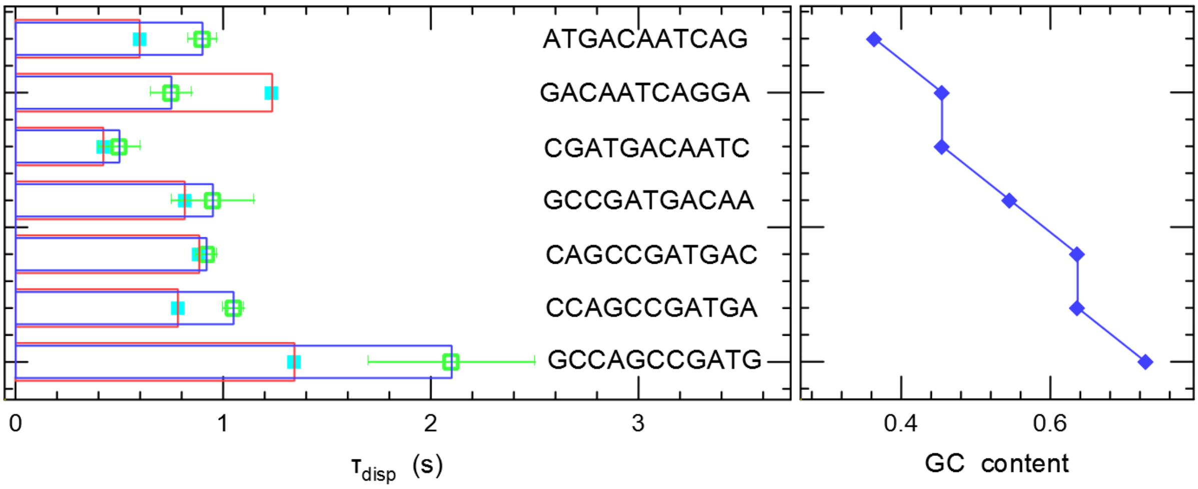


**Fig. S11** : The displacement time of 11nts oligonucleotides with different GC content at a testing force $F_{test}$=8.6 pN. Top: $<\tau_{disp}>$ for the 7 nts oligonucleotide, bottom: their GC content. The $<\tau_{disp}>$ values of 7 different oligonucleotides shown here do not vary simply with their GC content, i.e. for the melting temperature.

#### Salt Dependence

Using 1.2 kbps hairpin, 9nts oligomer ACAGCCAGC, at T=25°C, T4 buffer. We should replace "buffer dilution" by the salt concentration, the mean value of the two graphs are not really the same …

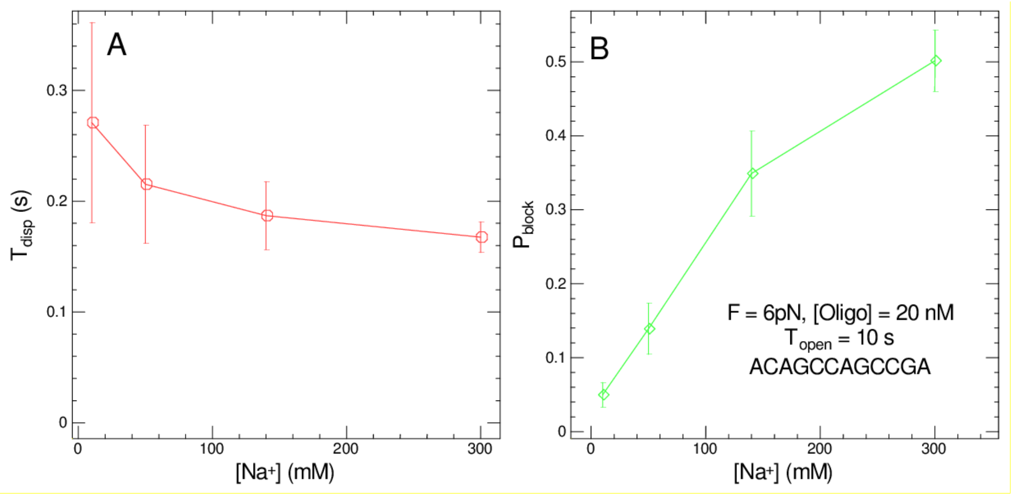


**Fig.S12**. Evolution of (A) of $<\tau_{disp}>$ with the ionic concentration and the oligonucleotide and (B) $P_{block}$ with the ionic concentration.

The oligonucleotide detachment starts from a dsDNA structure, where the Manning condensation is important, the hybridization is the encountering of two ssDNA molecules where the Manning condensation is weak. The salt entropy correction is related to the screening effect of salt ions surrounding the DNA molecules. This correction ought to be stronger for ssDNA and weaker for dsDNA, since detachment mostly occurs in dsDNA state the correction ought to be smaller.

#### Fork blocking assay with the 95 nt oligonucleotide

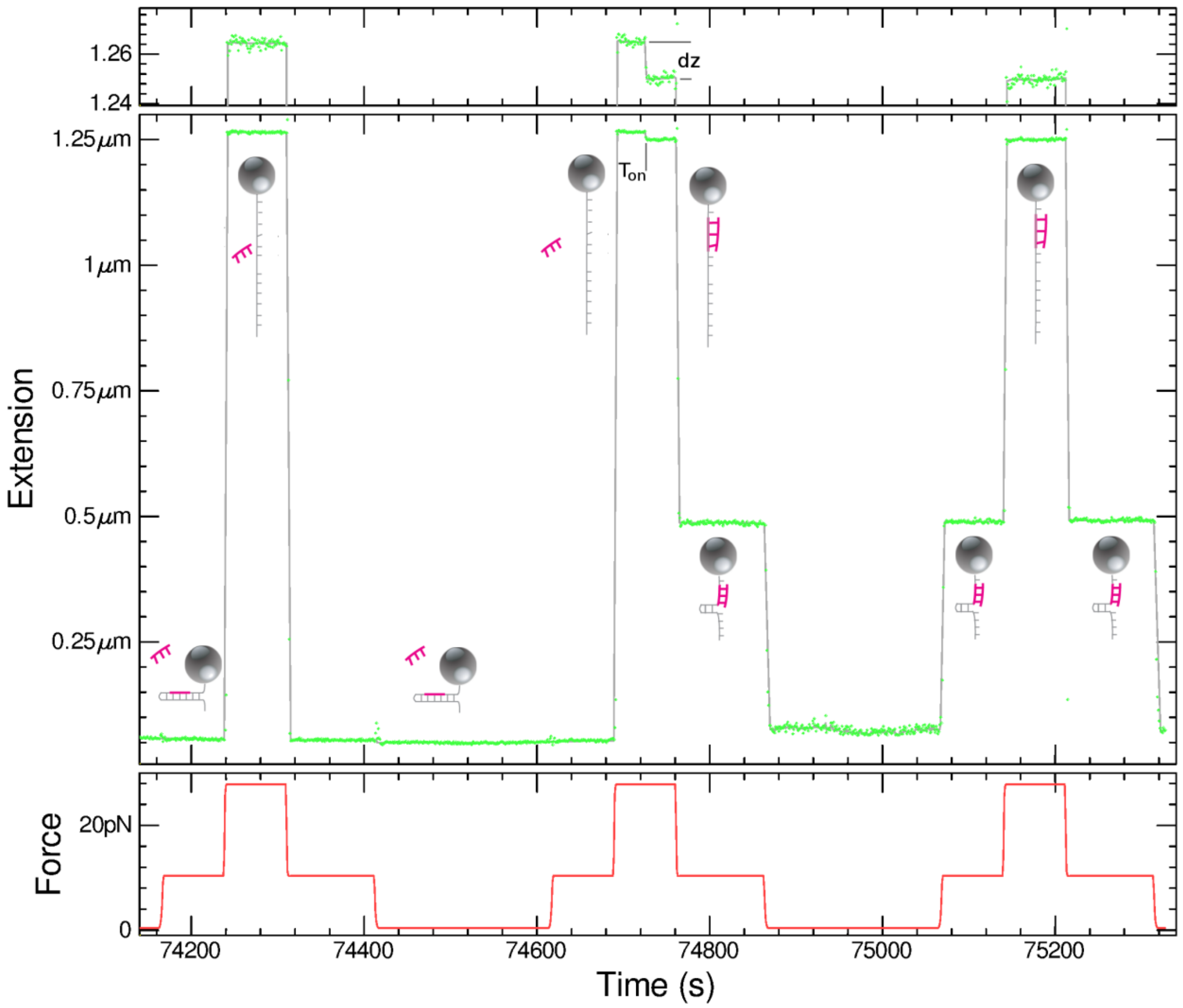


**Figure S13**: Fork blocking assay with the 95 nt oligonucleotide illustrating the detection of the oligonucleotide hybridization in the open phase. A 1.2 kb hairpin is subject to cycles where the force alternates between three values. A high force $F_{open}$ > 20 pN which unzips the hairpin, a medium force $F_{test}$ ~10 pN which reveals the hybridization state through the blocking of the hairpin at finite extension. A low force regime F ~0.5 pN which usually detach the hybridized oligonucleotide. The time course shown here displays three cycles, in the first one no oligonucleotide hybridizes as revealed by the hairpin extension which is basically zero except during $F_{open.}$ In the second cycle, an oligonucleotide hybridizes during the open phase, this is visible through the small change of extension dz occurring after Ton in the open phase. Moreover, when the force is decreased to $F_{test}$, the refolding is blocked. In the third cycle, the previously hybridized oligonucleotide is not expelled after the small force phase, this is evidenced by the blocking position observed when the force is raised at $F_{test}$ at t = 75075s. As the oligonucleotide is already hybridized, it remains during the open phase and blocks again the hairpin when the force is decreased to $F_{test}$ at t = 75220s. The top trace shows details of the extension in the open phase, with such a long oligonucleotide (95nts), the hybridization appears as an extension reduction of ~15 nm.

#### Pairing parameters

For DNA oligonucleotides we use Mfold 3.1 parameters. As we have measured that the off rate of the oligonucleotide does not depend on salt (see below) while the on-rate does depend, we have removed the salt correction in the entropy when computing off-rates. For the initiation energy and the AT penalty at oligonucleotide ends, we have used the virtual first and last dinucleotides where these dinucleotide free energy is corrected by half the initiation energy plus eventually the AT penalty. However, we have found that this reduction of binding energy appears too large for the end of the nucleotide that is competing with the fork, thus for that end, we have not applied this correction. The correction was applied on all free ends of the oligonucleotide that is one in the fork blocking configuration and two in the unpeeling one.

**Table (S3)**

| DNA/DNA | xA | | xT | | xC | | xG | |
|---|---|---|---|---|---|---|---|---|
| | *ΔΔH Kcal/mol* | *ΔΔS cal/mol* | *ΔΔH Kcal/mol* | *ΔΔS cal/mol* | *ΔΔH Kcal/mol* | *ΔΔS cal/mol* | *ΔΔH Kcal/mol* | *ΔΔS cal/mol* |
| Ax | -7.9 | -22.25 | -7.2 | -21.375 | -8.4 | -22.45 | -7.8 | -21.025 |
| Tx | -7.2 | -21.35 | -7.9 | -22.25 | -8.2 | -22.25 | -8.4 | -22.45 |
| Cx | -8.5 | -22.725 | -7.8 | -21.025 | -8.0 | -19.85 | -10.6 | -27.2 |
| Gx | -8.2 | -22.25 | -8.4 | -22.45 | -9.8 | -24.375 | -8.0 | -19.85 |
| | InGC | | 0.1 | -2.8 | InAT | | 2.3 | 4.1 |

| RNA/DNA | xA | | xT | | xC | | xG | |
|---|---|---|---|---|---|---|---|---|
| | *ΔΔH Kcal/mol* | *ΔΔS cal/mol* | *ΔΔH Kcal/mol* | *ΔΔS cal/mol* | *ΔΔH Kcal/mol* | *ΔΔS cal/mol* | *ΔΔH Kcal/mol* | *ΔΔS cal/mol* |
| Ax | -7.8 | -21.9 | -8.3 | -23.9 | -5.3 | -12.3 | -9.1 | -23.5 |
| Tx | -7.8 | -23.2 | -11.5 | -36.4 | -8.6 | -22.9 | -10.4 | -28.4 |
| Cx | -8.0 | -26.1 | -7.0 | -19.7 | -9.3 | -23.2 | -16.3 | -47.1 |
| Gx | -5.5 | -13.5 | -7.8 | -21.6 | -8.0 | -17.5 | -12.8 | -31.9 |
| | InGC | | 0.1 | -2.8 | InAT | | 2.3 | 4.1 |

| LNA/dna | xa | | xt | | xc | | xg | |
|---|---|---|---|---|---|---|---|---|
| | *ΔΔH Kcal/mol* | *ΔΔS cal/mol* | *ΔΔH Kcal/mol* | *ΔΔS cal/mol* | *ΔΔH Kcal/mol* | *ΔΔS cal/mol* | *ΔΔH Kcal/mol* | *ΔΔS cal/mol* |
| Ax | -7.193 | -19.723 | -4.918 | -12.943 | -7.269 | -17.336 | -7.536 | -18.387 |
| Tx | -7.246 | -19.738 | -6.372 | -16.902 | -6.307 | -15.515 | -10.040 | -25.744 |
| Cx | -7.451 | -18.380 | -7.092 | -16.825 | -5.904 | -11.904 | -9.815 | -23.491 |
| Gx | -5.036 | -11.656 | -8.612 | -23.327 | -10.160 | -24.651 | -10.844 | -25.580 |
| | InGC | | 0.1 | -2.8 | InAT | | 2.3 | 4.1 |

| dna/LNA | xA | | xT | | xC | | xG | |
|---|---|---|---|---|---|---|---|---|
| | *ΔΔH Kcal/mol* | *ΔΔS cal/mol* | *ΔΔH Kcal/mol* | *ΔΔS cal/mol* | *ΔΔH Kcal/mol* | *ΔΔS cal/mol* | *ΔΔH Kcal/mol* | *ΔΔS cal/mol* |
| ax | -6.908 | -18.135 | -5.384 | -13.537 | -5.510 | -11.824 | -9.000 | -22.826 |
| tx | -5.609 | -16.019 | -5.574 | -14.149 | -7.591 | -19.031 | -6.335 | -15.537 |
| cx | -7.142 | -18.333 | -9.471 | -25.070 | -5.937 | -12.335 | -10.876 | -27.918 |
| gx | -7.756 | -19.302 | -9.035 | -22.742 | -10.725 | -25.511 | -8.943 | -20.833 |
| | InGC | | 0.1 | -2.8 | InAT | | 2.3 | 4.1 |

| LNA/LNA | xA | | xT | | xC | | xG | |
|---|---|---|---|---|---|---|---|---|
| | *ΔΔH Kcal/mol* | *ΔΔS cal/mol* | *ΔΔH Kcal/mol* | *ΔΔS cal/mol* | *ΔΔH Kcal/mol* | *ΔΔS cal/mol* | *ΔΔH Kcal/mol* | *ΔΔS cal/mol* |
| Ax | -9.991 | -27.175 | -14.703 | -40.750 | -11.389 | -28.963 | -12.793 | -31.607 |
| Tx | -10.318 | -26.108 | -10.419 | -27.683 | -9.166 | -21.535 | -10.046 | -22.591 |
| Cx | -14.177 | -35.498 | -15.737 | -41.218 | -15.399 | -36.375 | -14.558 | -35.239 |
| Gx | -13.959 | -35.097 | -17.361 | -45.858 | -16.109 | -40.738 | -13.022 | -26.673 |
| | InGC | | 0.1 | -2.8 | InAT | | 2.3 | 4.1 |

Stability changes in RNA-DNA nearest neighbor pairing parameters for the oligonucleotide ACAGCCAGCCG using parameters at (1M NACl, 10 mM $Na_2$ $HPO_4$ and 1 mM $Na_2$ EDTA, pH 7.0) of (4). Note that the values given in the table are not extremely precise because they are estimated from the histograms in Fig.1 of (4). Moreover the ionic conditions are different from ours. The last base pair energy is not taken into account because we consider in the model that the oligonucleotide with fewer than 2 attached base pairs are unstable. The change of stability we fit from the data is 1.1 Kcal/mol and is to be compared with the sum of the pairing energies given in the table.

***Table S4**: ΔΔG Base pairing parameters from (4) for the RNA -DNA hybrid hairpin of 11 bases pairs*

| RNA/DNA | xA | | xT | | xC | | xG | |
|---|---|---|---|---|---|---|---|---|
| | *ΔΔH Kcal/mol* | *ΔΔS cal/mol* | *ΔΔH Kcal/mol* | *ΔΔS cal/mol* | *ΔΔH Kcal/mol* | *ΔΔS cal/mol* | *ΔΔH Kcal/mol* | *ΔΔS cal/mol* |
| Ax | -7.8 | -21.9 | -8.3 | -23.9 | -5.3 | -12.3 | -9.1 | -23.5 |
| Tx | -7.8 | -23.2 | -11.5 | -36.4 | -8.6 | -22.9 | -10.4 | -28.4 |
| Cx | -8.0 | -26.1 | -7.0 | -19.7 | -9.3 | -23.2 | -16.3 | -47.1 |
| Gx | -5.5 | -13.5 | -7.8 | -21.6 | -8.0 | -17.5 | -12.8 | -31.9 |
| | InGC | | 0.1 | -2.8 | InAT | | 2.3 | 4.1 |

In the table r refers to RNA and d to DNA.

#### Description of released programs:

**ToffOligo.py:** is a Python program that computes the displacement time of a DNA oligonucleotide for a given force and temperature. These parameters are provided by the users on the command line as options. The user can specify the oligonucleotide sequence using the option –O=XYZ or –-oligo=XYZ. By default the temperature is set to 25°C and the force to 8.5 pN. The user may modify these default settings using the options –T=value or –-temperature= value and –F=value or –force=value. For example, the command below displays the result displacement time of 1.1133 s:

python3 **ToffOligo.py** -F=8.6 -T=20 -O=ATGACAATCAG
For DNA oligo 5'-ATGACAATCAG-3' of 11 bp
at T 20C, F = 8.6 pN, matrix size 220

ATGACAATCAG 1.1133

With the option –h or –help, the option usage is reminded

Command line for the program

Simplest option the oligo alone and then describe all possible options :

oligo sequence, temperature, length, force, salt, DNA-DNA, DNA-LNA, DNA-RNA, apex, fork, mismatches only for DNA,
